# Cell states and neighborhoods in distinct clinical stages of primary and metastatic esophageal adenocarcinoma

**DOI:** 10.1101/2024.08.17.608386

**Authors:** Josephine Yates, Camille Mathey-Andrews, Jihye Park, Amanda Garza, Andréanne Gagné, Samantha Hoffman, Kevin Bi, Breanna Titchen, Connor Hennessey, Joshua Remland, Erin Shannon, Sabrina Camp, Siddhi Balamurali, Shweta Kiran Cavale, Zhixin Li, Akhouri Kishore Raghawan, Agnieszka Kraft, Genevieve Boland, Andrew J. Aguirre, Nilay S. Sethi, Valentina Boeva, Eliezer Van Allen

**Author notes:** these authors contributed equally (Josephine Yates, Camille Mathey-Andrews). these authors contributed equally (Valentina Boeva, Eliezer Van Allen).

## Abstract

Esophageal adenocarcinoma (EAC) is a highly lethal cancer of the upper gastrointestinal tract with rising incidence in western populations. To decipher EAC disease progression and therapeutic response, we performed multiomic analyses of a cohort of primary and metastatic EAC tumors, incorporating single-nuclei transcriptomic and chromatin accessibility sequencing, along with spatial profiling. We identified tumor microenvironmental features previously described to associate with therapy response. We identified five malignant cell programs, including undifferentiated, intermediate, differentiated, epithelial-to-mesenchymal transition, and cycling programs, which were associated with differential epigenetic plasticity and clinical outcomes, and for which we inferred candidate transcription factor regulons. Furthermore, we revealed diverse spatial localizations of malignant cells expressing their associated transcriptional programs and predicted their significant interactions with microenvironmental cell types. We validated our findings in three external single-cell RNA-seq and three bulk RNA-seq studies. Altogether, our findings advance the understanding of EAC heterogeneity, disease progression, and therapeutic response.

## Introduction

Esophageal adenocarcinoma (EAC) is believed to arise from Barrett’s esophagus, an uncommon metaplastic condition^1–7^. EAC is exceptionally lethal, with a 5-year survival rate of under 5% for patients with non-resectable disease or detectable metastases, representing over half of diagnosed patients^7,8^. The recalcitrant and heterogeneous response to treatment underscores the need to understand EAC progression at a cellular level and delineate malignant cell and tumor microenvironment (TME) heterogeneity in therapy-resistant and metastatic settings^4,9^.

While recent studies explored EAC at single-cell resolution to identify candidate immune and stromal cell types relevant to pathogenesis^9,10^, malignant cell states and their heterogeneity in EAC across disease stages — crucial for predicting disease progression, metastasis, and therapeutic response — remain largely undetermined^11,12^. Moreover, epigenetic heterogeneity, vital for understanding malignant cell plasticity^12^, as well as spatial relationships between distinct cell types and states, remain unexplored in EAC. Given recent advances of single-cell and spatial transcriptomics studies^13–15^, we hypothesized that joint inference of transcriptional, epigenetic, and spatial heterogeneity in EAC across disease stages, metastatic foci, and therapeutic exposures may provide novel insights into programs dictating lethal disease. Our analysis uncovered malignant cell programs and their spatial localizations and interactions with microenvironmental cell types that inform EAC disease progression and therapeutic resistance.

## Results

### Characterizing the transcriptional and chromatin accessibility landscape of primary and metastatic EAC

For our discovery cohort, we analyzed a total of 10 biopsies from therapy-naïve and therapy-exposed EAC patients using multiome sequencing (single-nuclei RNA sequencing [snRNA-seq] and single-nuclei ATAC sequencing [snATAC-seq]), and Visium spatial transcriptomics (ST) for a subset of 5 matched samples from 3 patients (Fig. 1a; Suppl. Fig. S1; Methods). Metastatic tumors were obtained from diverse anatomical sites, including three from the liver, one from the adrenal gland, and one from the peritoneum (Suppl. Fig. S1).

**Fig 1:**
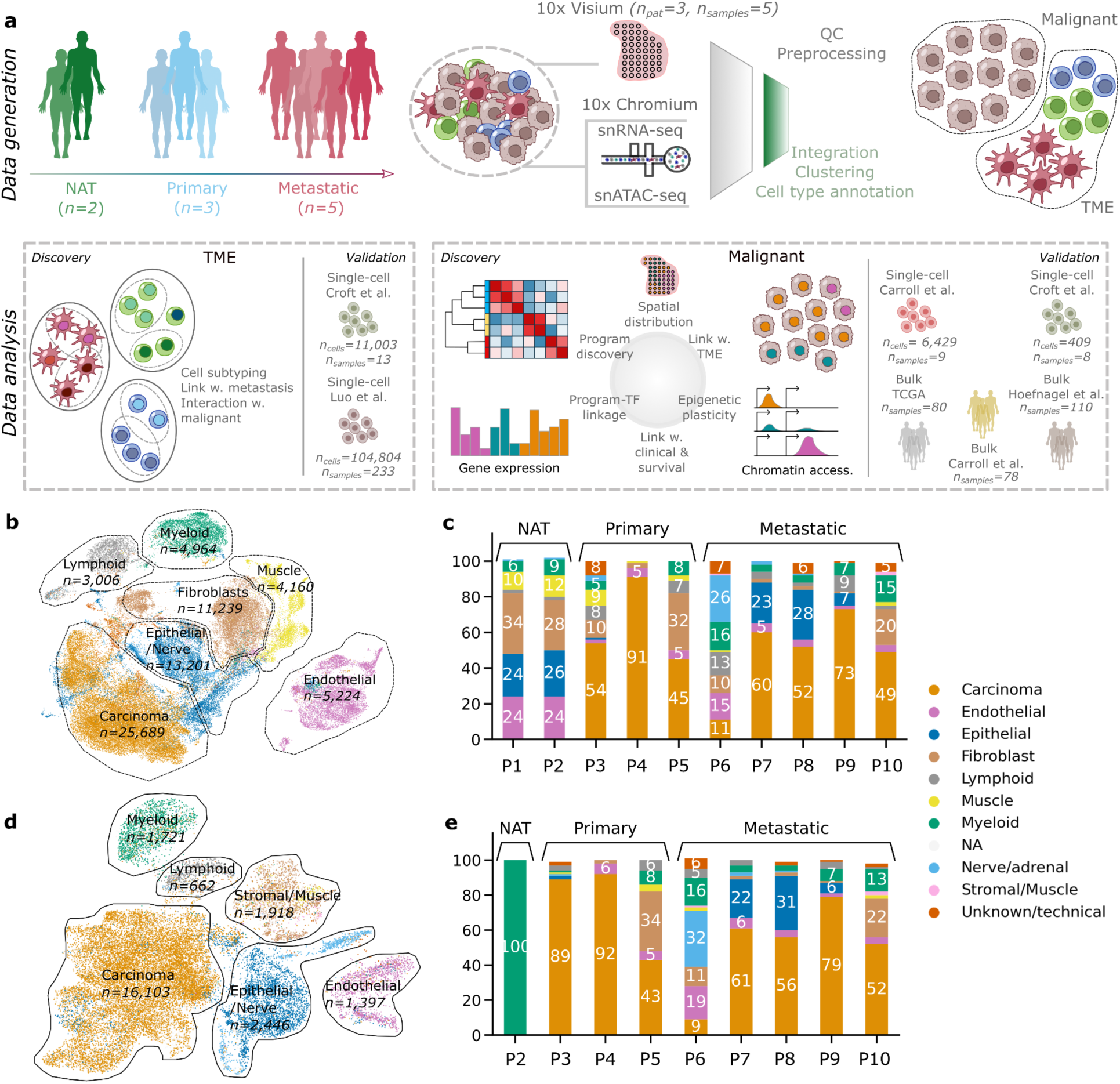
EAC primary and metastatic samples show a diverse landscape of TME and malignant cells in transcriptomic and epigenetic data. **a**, Schematic representation of the study workflow. Biopsies from 10 patients in our discovery cohort, including normal adjacent tissue (NAT), primary tissue, and metastatic samples, were subjected to single-nuclei RNA and ATAC sequencing using 10X Chromium technology. For a subset of these patients, matched primary and metastatic samples were sequenced with 10X Visium spatial transcriptomics (ST) technology. For single-nuclei data, cells were annotated by cell type and categorized into malignant and TME components. TME subtypes were linked to metastasis, with validation against an external pan-cancer fibroblast atlas^16^. The malignant cell components underwent analysis using consensus non-negative matrix factorization (cNMF) to uncover malignant programs, which were further characterized for transcriptional and epigenetic heterogeneity at a single-cell and spatial level and candidate master transcription factors. External validation was performed in two single-cell validation cohorts^9,10^, and associations with clinical and molecular characteristics, as well as survival, were assessed in three bulk validation cohorts^7,10,17^. b, Uniform Manifold Approximation and Projection (UMAP) representation of the full cohort in Harmony-corrected integrated transcriptomic data, with major cell type compartments labeled and cell counts indicated. c, Proportion of major cell types in each sample based on transcriptomic data, with percentages for compartments representing over 5% of the total sample composition. d, UMAP representation of the full cohort in Harmony-corrected integrated ATAC data, with cell type annotations transferred from the RNA annotations. “NA” denotes cells without paired associated RNA information. e, Proportion of major cell types in each sample based on ATAC data, with percentages for compartments representing over 5% of the total sample composition.

After preprocessing, we identified 72,552 high-quality cells with expression information for 21,444 genes within the snRNA-seq data and 33,966 cells with chromatin accessibility information for 311,978 genomic regions within the snATAC-seq data, represented for visualization purposes only in Harmony-corrected space (Methods; Fig. 1b-e; Suppl. Fig. S1). Seven major cellular compartments were delineated: carcinoma, epithelial/nerve, myeloid, muscle, fibroblast, and lymphoid, for which we uncovered various cell subtypes (Fig. 1b; Suppl. Fig. S2). Malignant cells represented an average of 54% of all cells across tumor samples (interquartile range, IQR: 48-63%); Fig. 1c).

### The EAC TME contains distinct macrophage and fibroblast populations

Although the response of EAC to immunotherapy can vary, recent studies have demonstrated that specific myeloid cell subtypes within EACs are associated with the effectiveness of immune checkpoint inhibitors (ICI)^9^. We found 5 distinct cell subtypes within the myeloid compartment, including two tumor-associated macrophage (TAM) populations (TAM1 and TAM2; Fig. 2a-b). TAM1 cells, exhibiting pro-inflammatory gene expression patterns^19–21^, were significantly enriched in tumor samples, whereas TAM2 cells, exhibiting characteristics of anti-inflammatory macrophages^22,23^, although present in tumor tissue, were differentially enriched in normal adjacent tissue (one-sample t-tests *p*<0.0001) (Fig. 2c)^18^.

**Fig 2:**
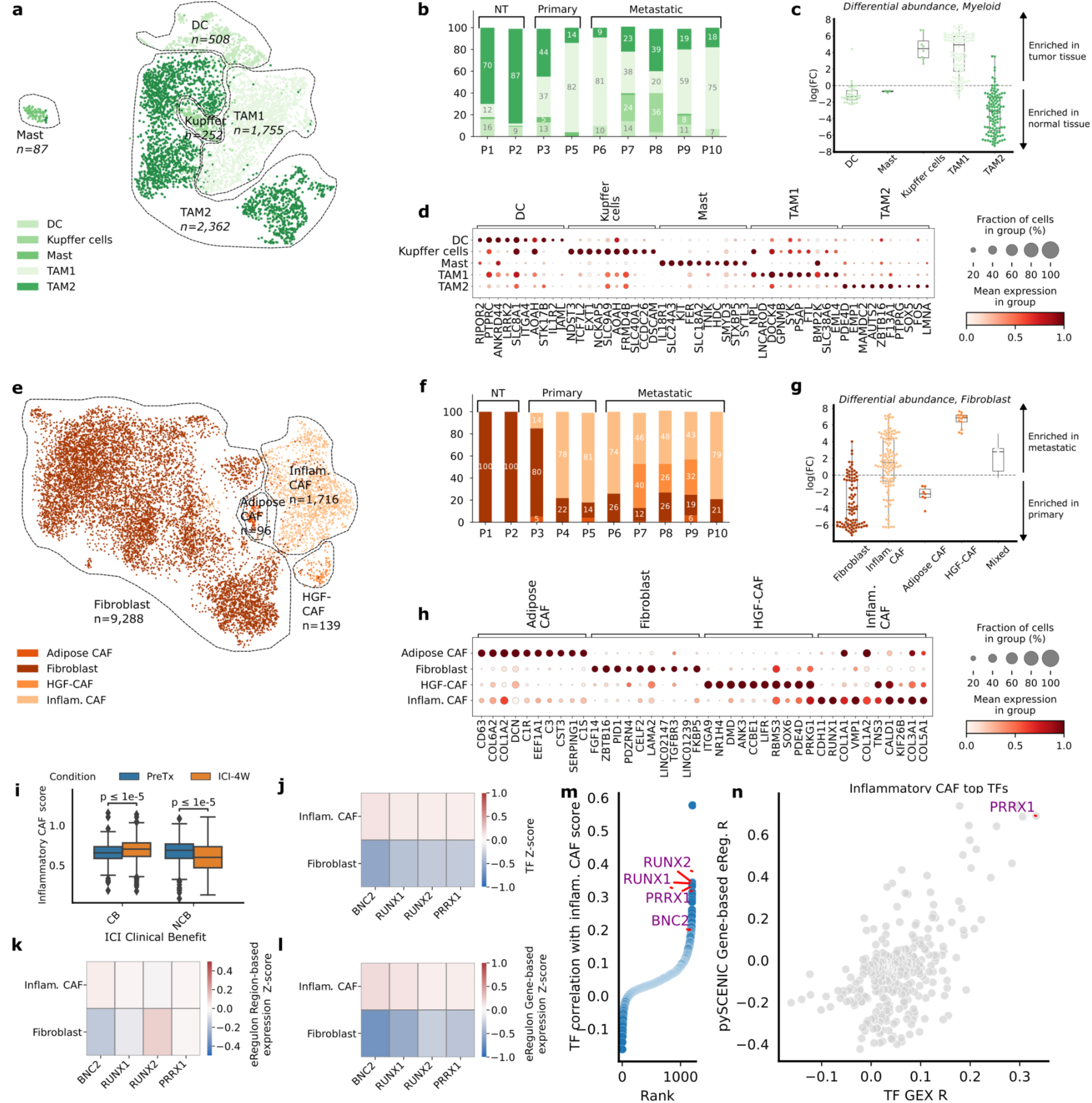
The EAC TME contains several pro– and anti-inflammatory populations of macrophages and RUNX1/RUNX2/PRRX1/BNC2-regulated inflammatory cancer-associated fibroblasts enriched in metastatic samples. **a**, Uniform Manifold Approximation and Projection (UMAP) representation of the myeloid compartment in Harmony-corrected integrated transcriptomic data, with annotated subtypes indicated. **b,** Proportion of myeloid subtypes per patient. **c,** Distribution of Milo^18^ fold-change scores between normal-adjacent and tumor samples for myeloid cells; Milo scores measure differential abundances of specific cell subtypes by assigning cells to overlapping neighborhoods in a *k*-nearest neighbor graph. **d,** Marker genes of annotated myeloid subtypes, with cells grouped by subtype and expression information provided. **e,** UMAP representation of the fibroblast compartment in Harmony-corrected integrated transcriptomic data, with annotated subtypes indicated. **f,** Proportion of fibroblast subtypes per patient. **g,** Distribution of Milo fold-change scores between metastatic and primary tumor samples for fibroblast subtypes, with labeling and exclusion criteria similar to (c). **h,** Marker genes of annotated fibroblast subtypes, with cells grouped by subtype and expression information provided. **i,** Distribution of the inflammatory cancer-associated fibroblast (CAF) score in the stromal compartment of the Carroll *et al*. ^9^ cohort, stratified by response to immune checkpoint inhibitor (ICI) therapy: clinical benefit (CB) and no clinical benefit (NCB). The inflammatory CAF program is scored on the entire cohort. Paired measurements of patients were made before treatment (PreTx) and after a 4-week ICI treatment window (ICI-4W). The distribution of the inflammatory CAF score is compared among the CB and NCB groups across PreTx and ICI-4W time points. Significance testing is conducted using a Mann-Whitney test to assess differences between the CB and NCB groups. **j-l,** Results for SCENIC+-derived transcription factor (TF) candidates for inflammatory fibroblasts, with cells grouped by subtype and Z-scores of TF expression (j), eRegulon gene-based expression (k), and eRegulon region-based expression (l) shown. **m,** TF gene expression correlation with inflammatory CAF score in the external pan-cancer fibroblast validation cohort of Luo *et al*. ^16^, with candidate TFs identified with the SCENIC+ analysis highlighted. **n,** Correlation of all available TFs’ gene expression and SCENIC-estimated gene-based eRegulon score with the inflammatory CAF score in the pan-cancer fibroblast atlas ^16^. Only PRRX1’s eRegulon activity, but not BNC2 and RUNX1/2, was estimated using SCENIC.

These TAM subpopulations resembled previously described populations in the pan-cancer tumor-infiltrating myeloid cell atlas^24^ and a study in EAC by Carroll *et al.*^9^ (Suppl. Fig. S3). Importantly, the TAM1 cells resembled the TAMs from the latter study, linked to higher monocyte content and selective ICI response, whereas TAM2 appeared similar to the M2 macrophages from the same study, linked to lower monocyte content and resistance to ICI.

Cancer-associated fibroblasts (CAFs) have also been previously implicated in tumor progression and therapy resistance^25,26^. We identified four distinct CAF populations in our cohort (Fig. 2e-f), including an inflammatory CAF population (iCAF) (expressing e.g., *CDH11, RUNX1, COL1A1*) enriched in metastatic EAC tumor samples and non-activated fibroblasts displaying relative abundance in primary EAC tumors (one-sample t-test *p*<0.0001) (Fig. 2g-h; Methods)^18^. These CAF populations were also consistently recovered in external pan-cancer and EAC-specific cohorts, encompassing a total of 246 tumor samples (Suppl. Fig. S4)^9,16^.

We next examined whether the presence of iCAFs correlated with selective ICI response in an external cohort^9^. Among the non-clinical benefit (non-CB) patient group (defined as the group of patients showcasing less than 12 months of progression-free survival), there was a significant decrease in inflammatory CAF gene signature scores following ICI treatment, consistent across patients, whereas a minimal increase, inconsistent across patients, in the inflammatory CAF score was observed pre– and post-ICI in the clinical benefit (CB) group (Fig. 2i; Suppl. Fig. S4).

Leveraging our paired snRNA-seq/ATAC-seq data, we used SCENIC+^27^ to identify candidate master transcription factor (mTF) regulons associated with the inflammatory CAF population^27^. RUNX1, RUNX2, PRRX1, and BNC2, previously implicated in various oncogenic processes^28–32^, were nominated as candidate mTFs of these cells (Fig. 2j-l) and further corroborated within the external pan-cancer CAF atlas^16^ (Fig. 2m-n). Overall, distinct macrophage and CAF cells associated with therapy response populate the microenvironment of both primary and metastatic EAC.

### Five malignant cell programs are identified across primary and metastatic EAC tumor samples

In contrast to TME investigations, tumor-intrinsic cellular programs relevant to progression, metastasis, and therapy resistance in EAC remain poorly understood^11,12,33^. To uncover unique gene activity programs operant among the EAC tumor compartment, we employed consensus non-negative matrix factorization (cNMF) and identified five cNMF programs consistently present across different patients (cNMF_1_ to cNMF_5_) (Fig. 3a-c; Methods). We conducted gene set enrichment analysis (GSEA) to assess enrichment of established biological pathways from MSigDB within the five cNMF malignant cell programs and compare the identified programs with the pan-cancer tumor cell programs from the pan-cancer study by Gavish *et al*.^33^ and the Barrett’s esophagus programs described by Nowicki-Osuch *et al.*^1^ (Fig. 3d, Suppl. Fig. S5).

**Fig 3:**
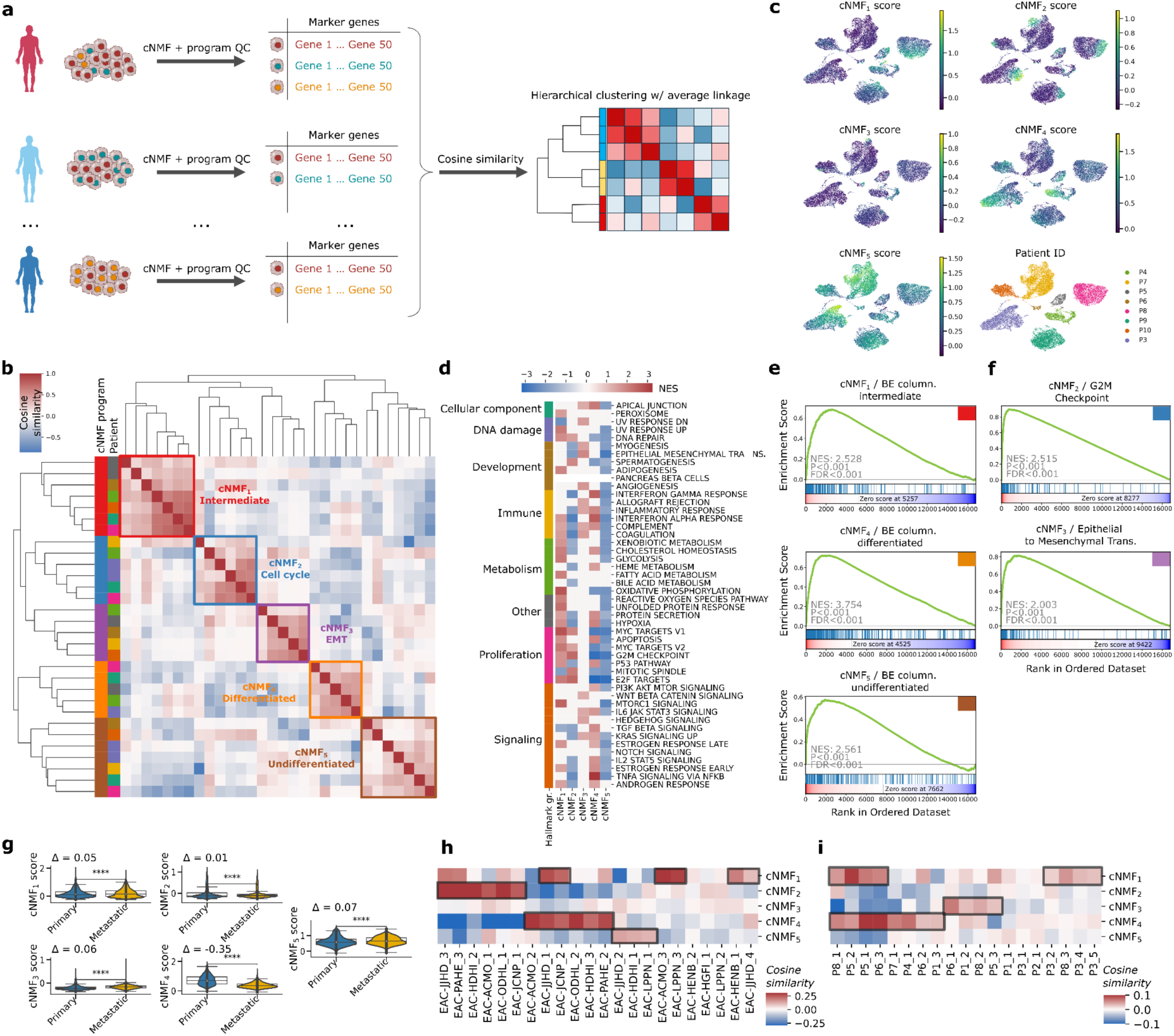
Five recurrent transcriptomic programs characterize EAC malignant cells with distinct RNA profiles. **a**, Illustration of the methodology employed for identifying transcriptomic programs. For each patient, consensus non-negative matrix factorization (cNMF) is performed on the malignant cell compartment, followed by manual filtration to retain high-quality programs characterized by gene weightings. Pairwise cosine similarity between programs across all patients is computed to cluster programs using hierarchical clustering with average linkage. **b,** Cosine similarity matrix representing the similarity between cNMF-derived programs across all samples, clustered using hierarchical clustering with average linkage. The five identified programs (cNMF_1_ through cNMF_5_) are delineated. **c,** UMAP representation of the malignant cell compartment using unintegrated transcriptomic data, colored according to their program score (cNMF_1_ through cNMF_5_) and sample ID. **d,** GSEA enrichment of the five programs in the 50 hallmarks of cancer, based on genes ranked according to their weight contribution to cNMF programs. Hallmarks are grouped according to category. Enrichments that did not reach significance (FDR=0.05) are blanked out. **e,** GSEA enrichment plots for selected programs described by Nowicki *et al*. in Barrett’s esophagus. **f,** GSEA enrichment plots for hallmarks G2M checkpoint in cNMF_2_ and Epithelial-to-Mesenchymal transition in cNMF_3_. **g,** Distribution of the five program scores in metastatic and primary samples. Significance is computed using the Mann-Whitney U test. The difference in median score is indicated as Δ. **h-i,** Cosine similarity between programs derived with cNMF in external datasets and cNMF_1_ through cNMF_5_ programs, derived in the Carroll *et al*. dataset (h) and in the Croft *et al*. dataset (i). The cosine similarity is computed between the cNMF-derived gene weights of programs for all patients in the external datasets and the median gene weight associated with each cNMF program derived in the discovery set.

cNMF_1_ resembled the intermediate columnar profile in Barrett’s esophagus^1^ (normalized enrichment score NES=2.5, FDR *q*<0.0001; Fig 3e), and showed enrichment in MYC targets, oxidative phosphorylation, and MTORC1 signaling pathways, akin to previously described Gavish *et al*. programs “EMT-III” and “Interferon/MHC-II (II)”. cNMF_2_ exhibited properties consistent with a cell cycling program (NES=2.5, FDR *q*<0.0001; Fig 3f), reminiscent of Gavish *et al*. program “Cell cycle G2/M”. cNMF_3_ resembled a classical EMT program (NES=2.0, FDR *q*<0.0001; Fig 3f), enriched in EMT and WNT beta-catenin pathways, aligned with the Gavish *et al*. program “EMT-I”. cNMF_4_ resembled the differentiated Barrett’s esophagus program (NES=3.8, FDR *q*<0.0001; Fig 3e)^1^, displayed enrichment in TNF, interferon-gamma, and interferon-alpha signaling, and appeared similar to the Gavish *et al*. “PDAC-classical”, “PDAC-related”, and “Epithelial senescence” programs. Finally, cNMF_5_ resembled the undifferentiated Barrett’s esophagus program (NES=2.6, FDR q<0.0001; Fig 3e).

Moreover, cNMF_4_ (differentiated esophagus program) was significantly enriched in malignant cells of primary EAC tumors (difference in median score between primary and metastatic malignant cells Δ=-0.35, p<0.0001), while cNMF_5_ (undifferentiated esophagus program) exhibited a slight enrichment in malignant cells of metastatic EAC samples (Δ=0.07, p<0.0001) (Fig. 3g).

To validate the robustness of the malignant cell cNMF programs uncovered in our study, we similarly performed cNMF on two external single-cell datasets sourced from Croft *et al*.^10^ and Carroll *et al*.^9^, across an aggregate of 6,838 malignant cells from 17 patient tumors. In the Carroll *et al*. dataset, we identified several programs consistent with cNMF_1_, cNMF_2_, cNMF_4_, and cNMF_5_; moreover, in the Croft *et al*. dataset, we observed enrichment of cNMF_1_, cNMF_3_, and cNMF_4_ programs, supporting the generalizability of the identified malignant cell programs across datasets (Fig. 3f-g, Suppl. Fig. S5).

### The five malignant cell programs displayed differential chromatin accessibility patterns and epigenetic plasticity

We next leveraged the paired snRNA-seq/ATAC-seq data to interrogate the connection between observed transcriptional programs and epigenetic diversity, aiming to decipher whether distinct EAC malignant cell programs correspond to specific chromatin accessibility patterns (Fig. 4a). We correlated the score of malignant cNMF programs with the normalized ATAC peak counts and identified significant associations between all cNMF programs and differentially accessible chromatin regions, denoted as cNMF-related peaks (Fig. 4b). We uncovered distinct chromatin accessibility patterns across cells representing cNMF programs (200 top-scoring cells, Methods), with several genes of interest displaying differential promoter accessibility, including *AKT2*^34^*, MKI67*^35^, *SPARC*^36,37^, *BHLHE41*^38,39^, and *ANXA11*^40^ (Fig. 4c). These variations in promoter and enhancer accessibility suggest a potential functional link between epigenetic alterations and evolution trajectories of tumor cells. Of note, cNMF_1_, cNMF_2_, and cNMF_3_ generally displayed less distinct chromatin accessibility profiles than cNMF_4_ and cNMF_5_.

**Fig 4:**
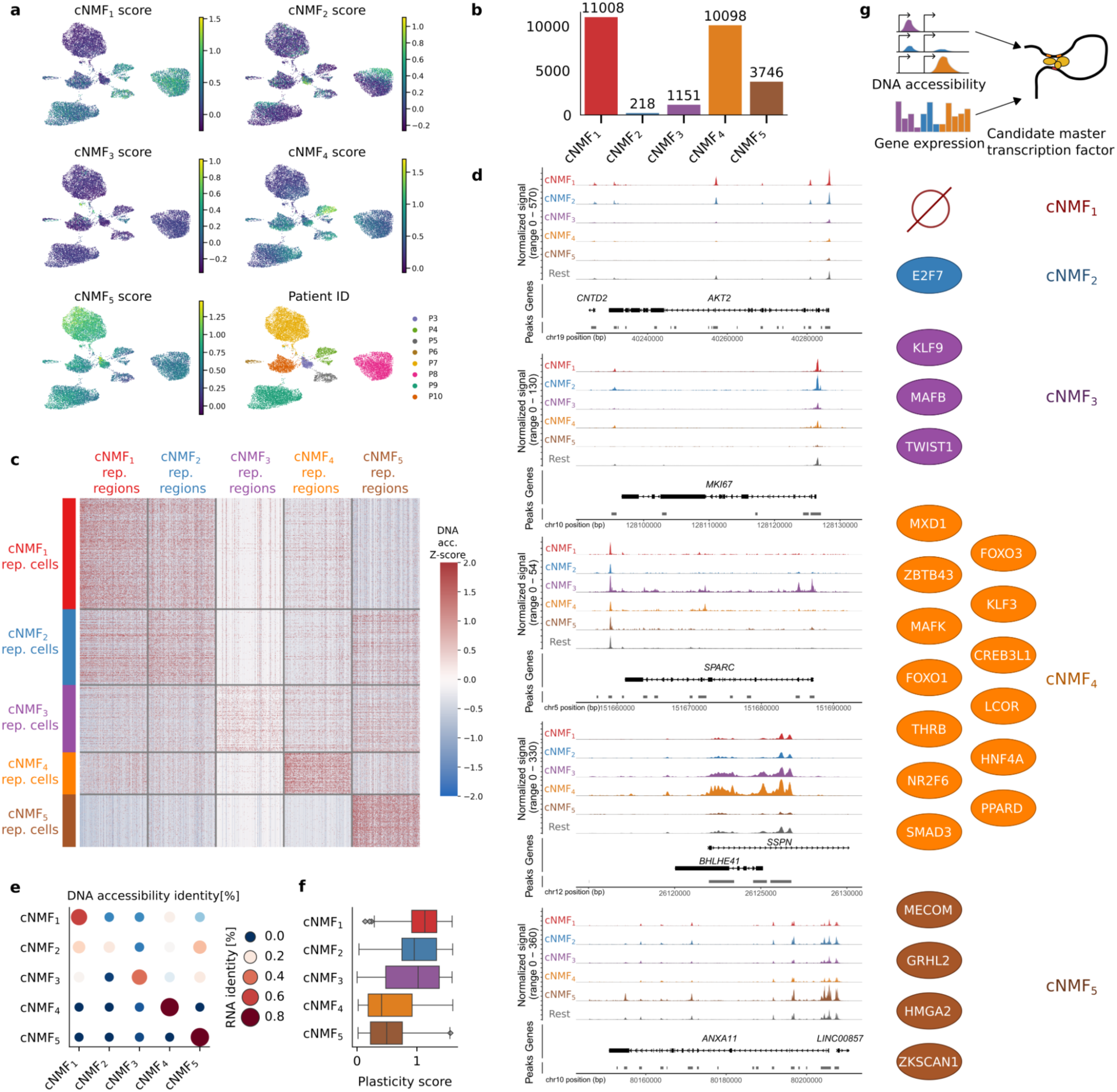
EAC malignant cell programs display unique ATAC profiles and epigenetic plasticity. **a**, UMAP representation of the malignant cell compartment using unintegrated snATAC-seq data, color-coded according to their cNMF gene signature score (cNMF_1_ through cNMF_5_) and sample ID. The program score is transferred from the RNA annotation. **b,** Number of open chromatin regions significantly correlated with each program (FDR<0.05, Pearson’s R>0.1). **c,** Heatmap illustrating chromatin accessibility in cNMF-associated regions for representative program cells. Cells are scored using cNMF signatures derived from RNA, with the top 5% unique cells in each score selected as representative cells. The top 200 regions with the higher correlation between chromatin accessibility and each program are represented. **d,** Chromatin accessibility of representative cNMF program cells for genes of interest. Genes are selected based on their association with the regions of the highest correlation between chromatin accessibility and gene signature scores of cNMF programs. Chromatin accessibility of promoters for *AKT2, MKI67, SPARC, BHLHE41,* and *ANXA11* is depicted for representative cells of cNMF_1_ through cNMF_5_ and all remaining carcinoma cells. **e,** The accuracy of classification of cells into cNMF programs using their chromatin accessibility profiles. Cells are scored by the average Z-score of chromatin accessibility of the top 200 cNMF-associated regions. The maximum score is used to classify cells into a chromatin accessibility identity; the percentage of cells from a gene expression identity classified into each chromatin accessibility identity is shown. **f,** Distribution of the epigenetic plasticity scores across representative cells of cNMF_1_ to cNMF_5_. Average Z-scores of ATAC accessibility vectors are transformed into a probability distribution using a softmax transformation with temperature, and the plasticity score is computed as the Shannon entropy over the resulting probability distribution. **g,** Representation of the candidate master transcription factors (mTFs) associated with programs consistent across datasets. We jointly model chromatin accessibility and gene expression to obtain candidate master transcription factors for each cNMF program in the discovery cohort that are subsequently validated in the two external validation cohorts. The identified mTFs consistent across datasets are represented.

Epigenetic plasticity, particularly the modulation of chromatin accessibility in malignant cells, is a recognized hallmark of cancer^41^. To determine if the identified cNMF programs exhibited chromatin states that facilitate transcriptional program diversity (epigenetic plasticity, as defined by Burdziak *et al.*^42^), we compared the paired transcriptional gene expression and chromatin accessibility profiles among malignant cells. Additionally, we analyzed the distribution of epigenetic plasticity scores within the malignant cells representing each cNMF program (Fig. 4d-f; Methods).

We assigned cells a gene expression (resp. chromatin accessibility) identity using the maximum signature score of signature genes (resp. cNMF-related peaks). Cells with strong cNMF_4_ and cNMF_5_ signature scores (within the top 5% of score distribution; Methods), representing differentiated and undifferentiated programs, respectively, exhibited mostly concordant transcriptional gene expression and chromatin accessibility identities, as well as low epigenetic plasticity, consistent with the hypothesized stable identity of these programs. Conversely, cells from the cell cycling program, cNMF_2_, displayed discordant expression of chromatin accessibility patterns characteristic of different programs along with high epigenetic plasticity^43,44^. cNMF_1_ also displayed high epigenetic plasticity, and certain malignant cells expressing the program had chromatin accessibility profiles that also associated with cNMF_4_ and cNMF_5_, consistent with the proposed intermediate nature of cNMF_1_ between the continuum represented by cNMF_5_ and cNMF_4_ programs (Fig. 3e).

Furthermore, cells within the EMT-like cNMF_3_ program displayed mixed chromatin accessibility identity and high epigenetic plasticity, consistent with previous observations of EMT state plasticity and its reversible nature^45,46^. Based on the snATAC-seq scores, *i.e.*, the average Z-score of normalized counts over cNMF-related peaks, we speculate that cNMF_3_ cells predominantly originate from the cNMF_1_ and cNMF_5_ pools rather than the cNMF_4_ pool, potentially suggesting that terminally differentiated EAC cells do not undergo EMT.

### Predicted transcription factor regulons of the malignant cell programs

To ascertain whether the identity of malignant cell cNMF programs was governed by a specific set of master transcription factors (mTFs), we next inferred the gene regulatory network underlying cell programs in our dataset leveraging the paired multiome data with SCENIC+^27^, and also evaluated these findings in the external Croft el al. and Carroll et al. datasets for reproducibility (Methods; Fig. 4g; Suppl. Fig. S6).

Candidate mTFs included E2F7^47^ for cNMF_2_; ZEB1^48,49^, TCF7L1^50^, and MAFB^51,52^ for cNMF_3_; FOXO1 and FOXO3^53^, MXD1^54,55^, LCOR^56,57^, CREB3L1^58–60^, MAFK^61^, PPARD^62,63^ and HNF4A^1,64–66^ (tumor suppressor TFs and/or associated with favorable prognosis) for cNMF_4_; and MECOM^67,68^ and HMGA2^69,70^ for cNMF_5_. Notably, no mTF was robustly identified across datasets for cNMF_1_. We therefore identified a set of candidate mTFs reproducibly associated with each malignant cell program except cNMF_1_ in three independent datasets (summarized in Fig. 4g). Lastly, the expression of genes coding for candidate mTFs identified for cNMF4 and cNMF5 was analyzed along the axis of expression of these two hypothesized opposing programs by ranking cells according to their relative cNMF_4_ to cNMF_5_ expression. The mTFs showed a consistent positive and negative gradient of expression along the cNMF_5_ to cNMF_4_ axis (Suppl. Fig. S6), supporting their role in orchestrating these program expressions.

### Malignant and TME cell programs in EAC display differential spatial enrichment in defined tumor regions

To assess whether the malignant cell programs identified in the snRNA-seq data exhibit spatial heterogeneity within individual EAC tumor samples, we performed Visium spatial transcriptomics (ST) on additional EAC tissue from matched patients (Methods). We categorized ST spots into pure tumor regions, mixed regions containing both malignant cells and TME cells in similar proportions, or regions of normal tissue, using CNV (copy number variation) profiles inferred from the spatial data (Methods, Fig. 5a-b, Suppl. Fig. S7). Deconvoluted ST spot cell type proportions and gene expression^71^ broadly agreed with CNV assignments (Suppl. Fig. S7-8).We scored the five malignant cell cNMF programs based on the corrected, deconvoluted carcinoma-specific gene expression matrix and found that they displayed distinct spatial distributions within the EAC tumor samples (Fig. 5a-b, Suppl. Fig. S7).

**Fig 5:**
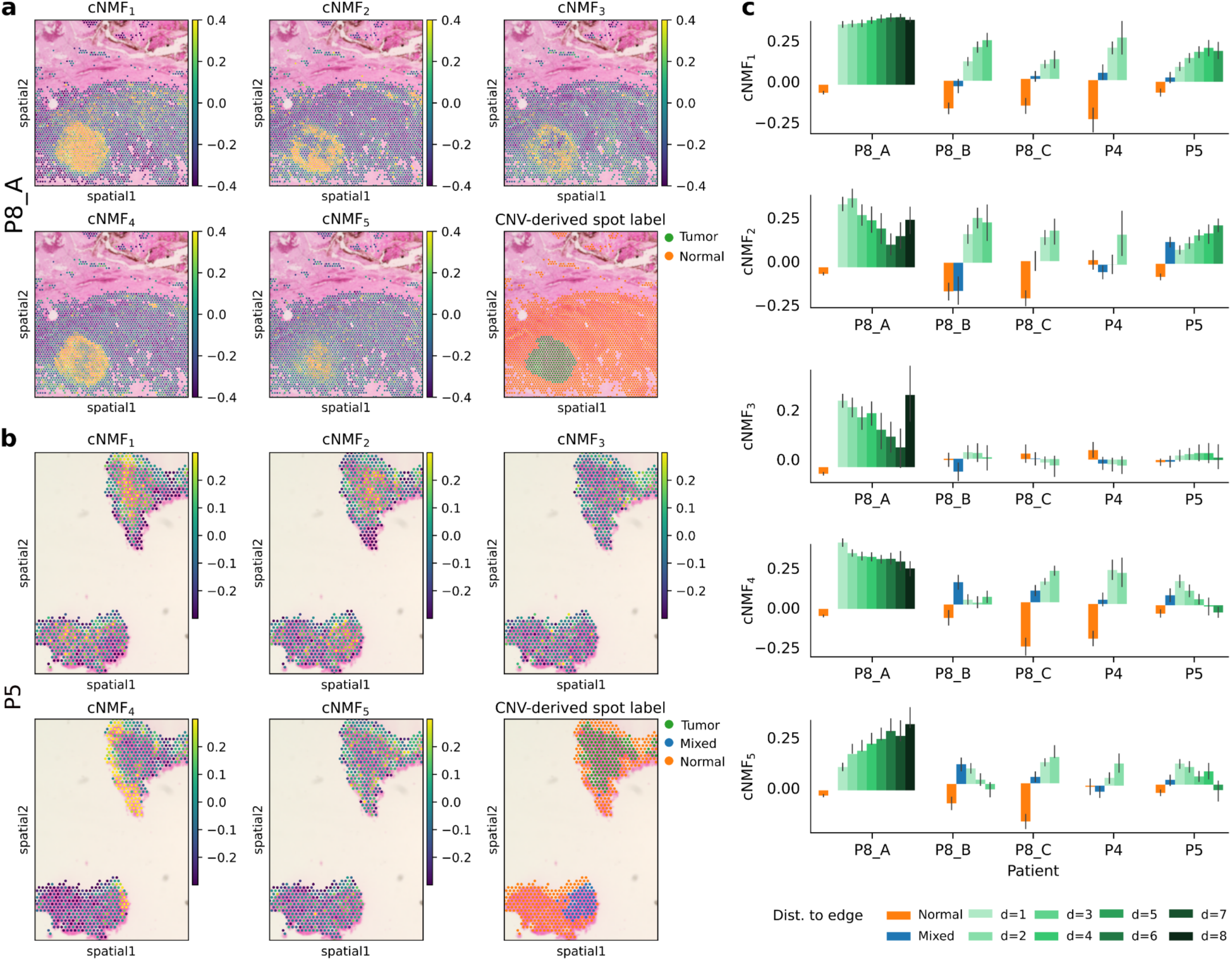
Single-nuclei derived transcriptional programs highlight different spatial regions of EAC tumors. **a-b**, Spatial transcriptomics (ST) slides of **a,** P8 primary tumor A and **b,** P5 primary tumor, colored according to cNMF program score and the CNV-derived label. For each spot, we infer the CNV profile with inferCNV and assign spots to tumor, mixed, and normal status. cNMF scores are computed as the average Z-score of signature genes using the deconvolved carcinoma-specific gene expression profile of spots derived with Cell2Location. **c**, Average cNMF score according to the position of the spots compared to the tumor leading edge. For each tumor spot, we compute the distance to the edge as the shortest path to a normal or mixed spot. The distribution of cNMF scores with standard error is represented for normal spots, mixed spots, and spots of a certain distance to the edge.

Specifically, in most samples, cNMF_1_ and cNMF_2_ were predominantly expressed in the tumor core, characterized by higher distances from the periphery; in contrast, cNMF_4_ was mainly expressed at the tumor periphery (Fig. 5c). cNMF_5_’s spatial location varied, while cNMF_3_, less frequently detected in snRNA-seq, was expressed in only three samples (P8_A, P8_B, and P5) and displayed dispersed spatial enrichment across the tumor (Fig. 5a-b, Suppl. Fig. S7). Thus, malignant cell programs exhibited reproducible and distinct spatial distributions within EAC tumors, although this association would require further validation through additional immunofluorescence staining.

### EAC malignant cell programs correlate with clinical characteristics, differential patient prognosis, and differential predicted drug sensitivity

We then sought to determine whether any of the identified malignant cell cNMF programs were associated with distinct clinical prognostic stages and therapeutically relevant states. By projecting these programs into the primary EAC TCGA cohort, we observed that cNMF_4_ was significantly linked with lower T and N stages, whereas cNMF_3_ exhibited a moderate association with higher T stages, consistent with its EMT-like nature^72^ (Fig. 6a-b). Other programs did not display significant associations with these clinical stages, and no malignant cell program showed significant associations with M staging (Suppl. Fig. S9).

**Fig 6:**
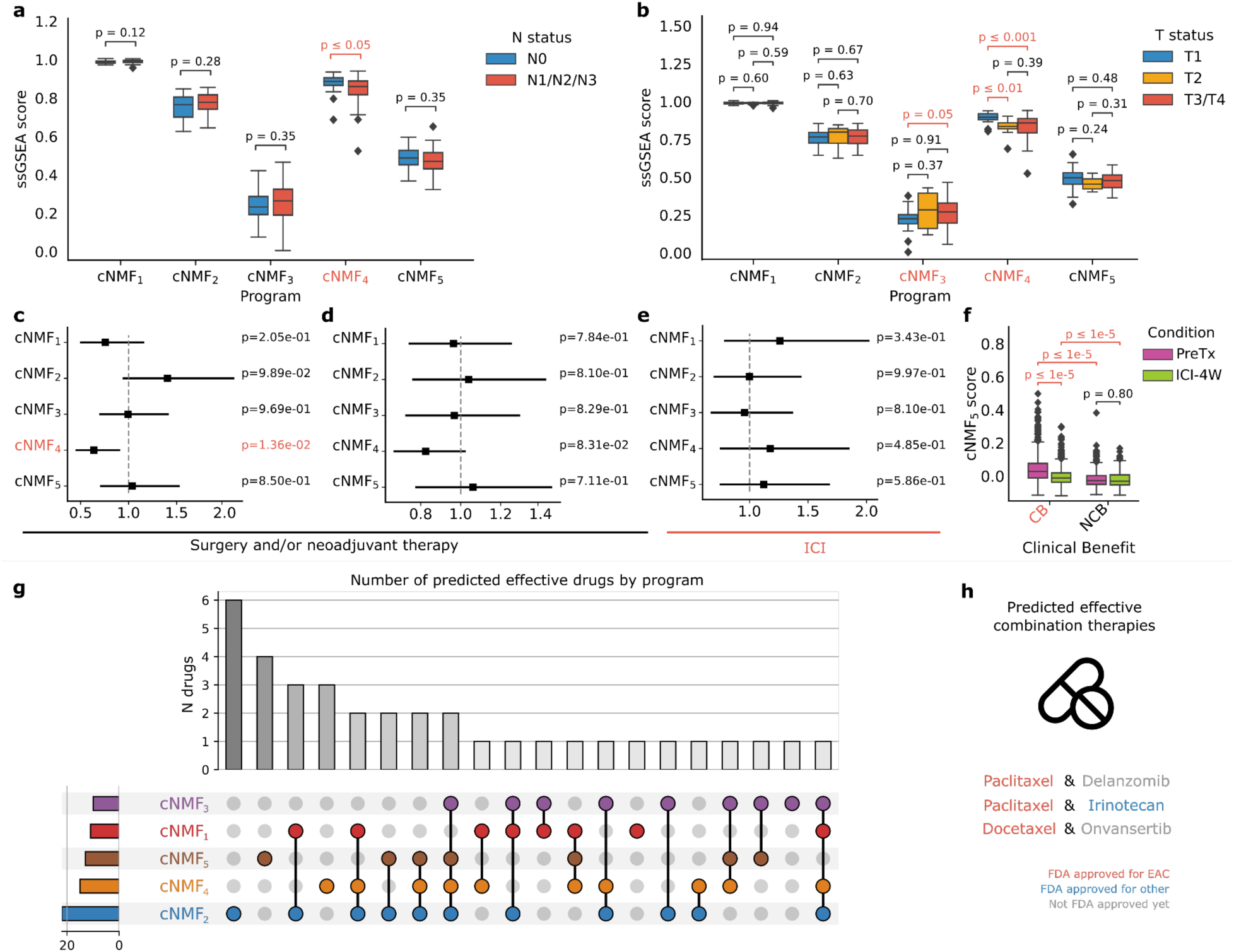
Discovered malignant programs have different clinical characteristics and predicted drug sensitivity. **a-b**, Link between uncovered programs and **a,** N stage, i.e., proxy of the number of nearby lymph nodes that have cancer, and **b,** T stage, i.e., size and extent of the main tumor in the TCGA bulk cohort ^7^. Patients are scored using single-sample Gene Set Enrichment Analysis (ssGSEA) with a cancer-specific gene signature. Statistical testing is performed using the Mann-Whitney U test. **c-e,** Hazard ratio associated with scores in bulk validation cohorts of **c,** TCGA, **d,** Hoefnagel *et al*. ^17^, **e,** and Carroll et al. ^9^. Cox proportional hazard univariate models are employed using disease-specific survival for TCGA and overall survival for Hoefnagel *et al*. and Carroll *et al*. **f**, Distribution of the cNMF_5_ score in the malignant cell compartment of the Carroll *et al*. cohort ^9^, stratified by response to immune checkpoint inhibitor (ICI) therapy: clinical benefit (CB) and no clinical benefit (NCB). The cNMF_5_ program is scored on the full cohort. Paired measurements of patients were made before treatment (PreTx) and after a 4-week ICI treatment window (ICI-4W). The distribution of the cNMF_5_ score is compared among the CB and NCB groups across PreTx and ICI-4W time points. Significance testing is conducted using a Mann-Whitney U test. **g,** Predicted drug sensitivity by program. scTherapy is used to infer to which drugs may exhibit activity in specific tumor programs. The upset plot represents the total number of drugs predicted to target a specific program on the left, as well as the size of the intersection represented on the middle panel on the top. **h,** Selected predicted candidate combination therapies that could target all five programs at a time are represented.

We then investigated the relationship between the malignant cell programs and patient survival using a univariate Cox proportional hazard model in two external bulk EAC patient cohorts treated with conventional therapies, namely surgery and neoadjuvant chemotherapy (TCGA^7^ and Hoefnagel et al.^17^), and one external EAC patient cohort treated with ICI (Carroll et al.^9^) (Methods). Higher cNMF_4_ scores were predictive of improved patient survival in the first two patient cohorts exposed to conventional therapies (p=0.01 and p=0.08 resp.) but not in the third patient cohort exposed to ICI (p=0.49) (Fig. 6c-e). The association of cNMF_4_ with less aggressive clinical features in this context is consistent with other program-specific features previously shown (*i.e.*, enrichment in primary tumors, differentiated transcriptional profile, and link to TFs associated with improved patient prognosis).

Finally, we investigated whether the cNMF programs displayed differential enrichment in therapy exposure categories of the external EAC patient cohort treated with ICIs. We assessed the distribution shift of the cNMF_5_ gene signature score in Carroll *et al.* single-cell data and observed the score was high in patients experiencing a clinical benefit (CB) to ICI both pre– and post-ICI exposure compared to non-CB patients (Mann Whitney U *p*<1e-5, Fig. 6f). In addition, the cNMF_5_ program gene signature score was significantly lower post-ICI exposure only in patients experiencing a CB (Fig. 6f). These patterns were consistent on a per-individual patient sample basis (Suppl. Fig. S9).

To assess whether the identified programs exhibit differential sensitivity to existing therapies, we used scTherapy^73^ to predict drug sensitivity for each program. Our analysis revealed a broad range of predicted responses, with cNMF2 showing sensitivity to the highest number of drugs, while cNMF3 displayed the lowest sensitivity (Fig. 6g). Notably, no single drug was predicted to target all five programs simultaneously; however, Delanzomib, a next-generation proteasome inhibitor^74^, Romidepsin, a HDAC inhibitor^75^, and SN-38, the active metabolite of irinotecan, an FDA-approved chemotherapy previously shown to have potential in EAC^76^, were the only agents predicted to affect four out of five programs.

Leveraging these predictions, we identified potential combination therapies by pairing drugs that together target all five cNMF programs (Fig. 6h). Among these combinations, some included FDA-approved chemotherapy agents for EAC, such as paclitaxel and docetaxel. Specifically, the combination of paclitaxel with delanzomib^74^ or with irinotecan^76^ was predicted to effectively target all five programs. Additionally, the combination of docetaxel and onvansertib, a drug recently granted FDA fast-track designation for metastatic colorectal cancer^77^, also emerged as a promising therapeutic strategy.

The cNMF programs identified in our study exhibited distinct associations with patient survival, therapy exposure status in external EAC cohorts, and predicted drug sensitivity. Notably, no single agent was predicted to target all five programs simultaneously, highlighting potential resistance mechanisms to standard chemotherapy. Furthermore, we suggest rational combination strategies that could be explored in preclinical models to overcome therapy resistance, offering a path toward more effective treatment approaches.

### Co-occurring groups of TME cells are linked with malignant cell programs

Lastly, given our findings of the key roles of tumor cell programs in EAC, we sought to understand whether and how the malignant cells interacted with specific TME cells (including the key myeloid and CAF populations described above). We first conducted an analysis of ecotypes, *i.e*., co-occurring abundance of tumor immune and stromal microenvironment cells, as measured in deconvolved data, in EAC^79^. Leveraging the external TCGA, Hoefnagel *et al*., and Carroll *et al*. EAC patient cohorts (n = 268 patients), we identified two major ecotypes: ‘immune-desert’ (predominantly comprising malignant cells and endothelial cells) and ‘immune-activated’ (featuring a mixture of myeloid, lymphoid, and stromal cells; Fig. 7a; Suppl. Fig. 9; Methods). A high cNMF_3_ gene signature score in deconvoluted samples was significantly associated with the immune-activated ecotype across all studies, in line with the described interaction of malignant cells with stromal and myeloid components to initiate EMT^80–82^ (Fig. 7a, Suppl. Fig. 9). Conversely, cNMF_5_ exhibited a significantly lower gene signature score in the immune-activated ecotype in two out of the three external cohorts (Fig. 7a, Suppl. Fig. 9).

**Fig 7:**
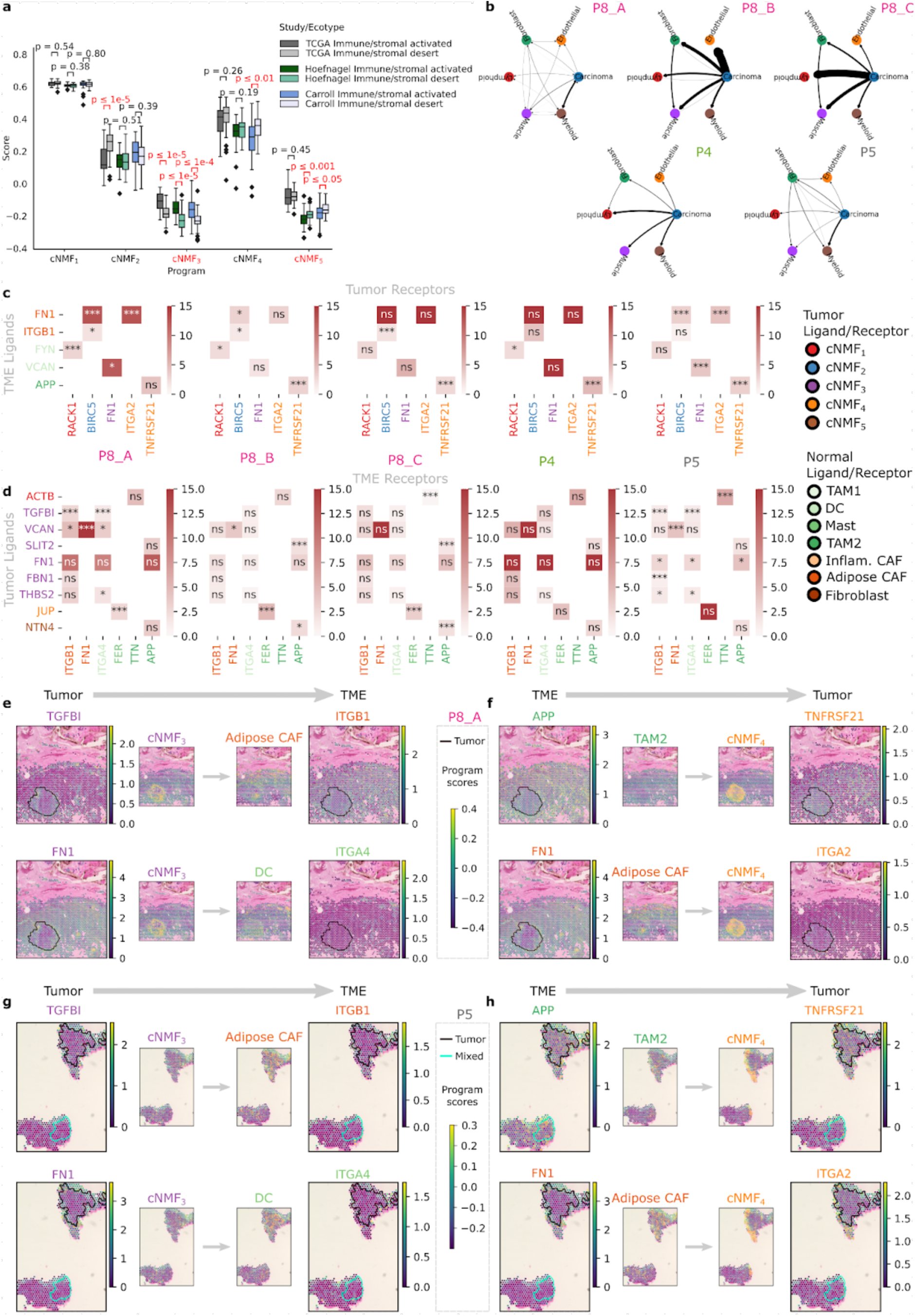
Uncovered malignant programs show associations with clinical and molecular characteristics, prognosis, and distinct ecotypes. **a**, Ecotype analysis of the data from the TCGA, Hoefnagel *et al*., and Carroll *et al*. cohorts deconvolved by BayesPrism. Distribution of cNMF scores in the two uncovered ecotypes, for each study. Statistical testing is performed using the Mann-Whitney U test. **b,** Estimated strength of interaction between cell types in spatial transcriptomics (ST) data. Using the NCEM method on Cell2Location-deconvolved data, we estimate in a spatially constrained manner the strength of interaction between cells from the 6 major compartments identified in the discovery cohort, represented for samples P8_A, P8_B, P8_C, P4, and P5. **c-d,** Significant ligand-receptor interactions uncovered with CellPhoneDB’s^78^ Squidpy implementation, LIGREC, for **c,** TME to tumor interactions and **d,** tumor to TME interactions. CellPhoneDB is run for each sample on spots near the edge of the tumor, defined as tumor spots (resp. normal spots) with a distance to the edge of less than 2. Only significant interactions (FDR *p*<0.1), for which the ligand/receptor is part of the signature genes of the cNMF programs/TME subtypes are represented. The ligand/receptor is colored according to which signature it belongs to. The hue encodes the CellPhoneDB mean of the ligand receptor pair; the level of significance is annotated for each existing interaction. ns: FDR p>0.1; *: 0.1≤p<0.01; **: 0.01≤p<0.001; ***: p≤0.001. **e-h,** Significant ligand-receptor interactions between **e,** P8 sample A tumor and TME components, **f,** P8 sample A TME and tumor components, **g,** P5 tumor and TME components, and **h,** P5 TME and tumor components. Each panel represents the log1p expression in spots. The ligand (resp. receptor) is colored according to the program whose signature genes it belongs to. Smaller panels represent the score distribution of the corresponding cNMF or TME component scores.

To further investigate significant interactions between malignant cells expressing differential cNMF program activity scores and TME cells, we predicted signaling interactions in our ST data (Methods)^83^. From this analysis, we observed that malignant cells had predicted signaling interactions with all TME compartments, but the strongest (i.e., highest impact on gene expression as predicted by NCEM; Methods) interactions were with myeloid and lymphoid cells (Methods; Fig. 7b)^9^.

Using computational cell-cell interaction prediction, we uncovered numerous candidate ligand-receptor interactions between malignant cells and myeloid and fibroblast subtypes (Methods; Fig. 7c-h; Suppl. Fig. 9)^78^. For example, we identified interactions between malignant cells with high cNMF_3_ activity scores and adipose CAFs, a small subpopulation of CAFs that co-occur with iCAFs and are predicted to be immunomodulatory^16^, through FN1 and ITGB1 and TGFBI and ITGA2^84^ that further support the association of cNMF_3_ and the immune-activated ecotype in external bulk datasets. We also found significant ligand-receptor interactions between cells displaying markers of TAM2 and adipose CAFs and malignant cells expressing cNMF_4_, potentially driven by the peripheral localization of cNMF_4_ discussed above.

Overall, these findings highlight the complex communication within the EAC TME and its spatial dependency, which could significantly influence EAC progression and treatment response.

## Discussion

Despite progress in dissecting EAC and Barrett’s esophagus biology, as well as relating biological programs to selective therapeutic response across treatment modalities, the complexities of its malignant cell compartment, epigenetic variations, and disease progression remain poorly understood. Leveraging a multi-modal profiling strategy across primary and metastatic EAC samples, our study unveiled considerable heterogeneity within and between tumors. In addition to identifying previously described myeloid and stromal compartments, this study is the first to define EAC malignant cell heterogeneity across primary and metastatic sites in distinct clinical settings across transcriptomic, chromatin accessibility, and spatial dimensions. We identified five major malignant cell programs, shared across patients in our study and external EAC patient cohorts, that possessed distinct chromatin accessibility profiles and spatial distributions. Among the programs identified, cNMF_5_, cNMF_1_, and cNMF_4_ delineated a continuum from undifferentiated to differentiated programs, mirroring a trajectory observed in Barrett’s esophagus^1^, cNMF_2_ represented a cell cycling program, and cNMF_3_ emerged as a rarer EMT-associated program (Fig. 8). Furthermore, we identified candidate transcription factors for various programs and a concordance between transcriptional programs and estimated epigenetic plasticity, contributing to the growing evidence emphasizing the significance of epigenetic plasticity as a facilitator of cancer progression and metastasis through increased heterogeneity^85–89^. We highlighted differential spatial distribution of malignant cell program gene signature scores and the tumor ecosystem’s complexity. Finally, we identified recurrent interactions between cells with high expression of cNMF programs and TME cell types, which could in turn influence therapy response, notably to ICI or targeted therapies that have shown strong dependency to TME cells and malignant cell heterogeneity^90–93^.

**Fig 8:**
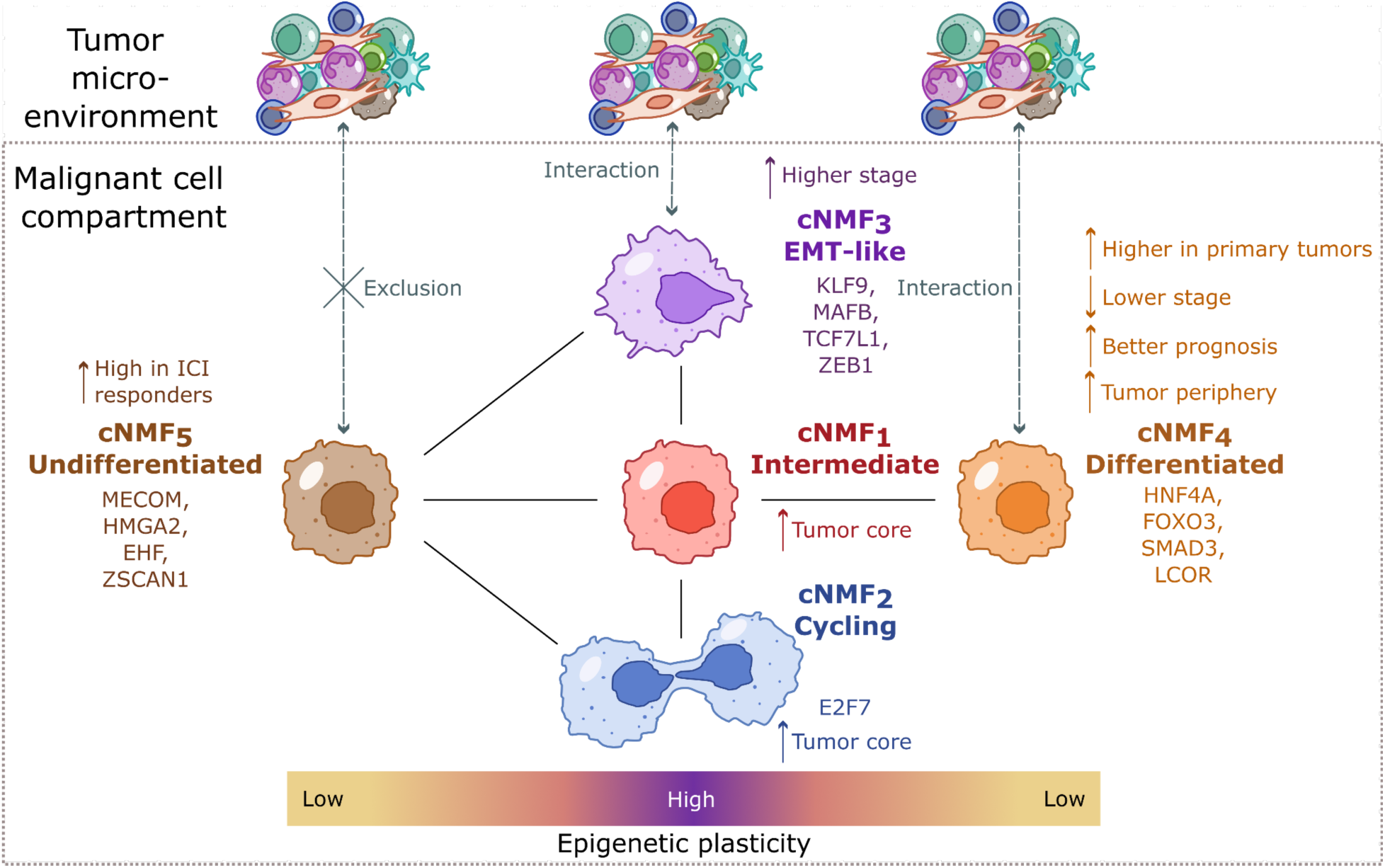
Summary of Key Findings in the Malignant Cell Compartment of Esophageal Adenocarcinoma (EAC). Five distinct malignant programs were identified, characterized by unique RNA and ATAC accessibility profiles. Among these, cNMF_5_ and cNMF_4_ represented two stable opposed programs: cNMF_5_ resembled an undifferentiated program, while cNMF_4_ exhibited characteristics of a differentiated program. cNMF_1_ displayed features of an intermediate program between cNMF_5_ and cNMF_4_. Conversely, cNMF_3_ manifested as a rare epithelial-to-mesenchymal transition (EMT)-like program, and cNMF_2_ represented a cell cycling program. ATAC accessibility profiles suggested potential transitions between these programs. Specifically, cNMF_3_ appeared epigenetically similar to cNMF_5_ and cNMF_1_ but distinct from cNMF_4_, and cNMF_2_ exhibited similarities with cNMF_5_ and cNMF_1_ but not cNMF_4_. The two hypothesized stable programs, cNMF_4_ and cNMF_5_, displayed lower epigenetic plasticity compared to the other programs. Candidate master transcription factors (mTFs) were identified for each transcriptional program. Furthermore, cNMF_5_ was associated with differential response to immune checkpoint inhibitor (ICI) therapy, while cNMF_4_ showed associations with lower T and N stages and better prognosis following surgery and/or neoadjuvant therapy treatment. In contrast, cNMF_3_ exhibited a slight enrichment in higher T stages. cNMF_1_ and cNMF_2_ were preferentially located at the tumor core, while cNMF_4_ was preferentially located at the tumor periphery. Lastly, cNMF_3_ and cNMF_4_ interacted significantly with the TME while cNMF_5_ was associated with immune exclusion.

There are several limitations in our study. Firstly, the discovery cohort comprised 10 samples, with 8 being tumor samples, hindering direct linkage between proportions of TME cells and clinical characteristics with malignant cell composition. Consequently, we mostly depended on bulk validation cohorts to elucidate these associations. Second, sampling and processing biases may affect differential abundance testing and limit the interpretability of the results. Notably, metastatic tumors were obtained from diverse anatomical sites, meaning that observed TME composition differences may reflect site-specific characteristics rather than metastatic status alone. Third, one of the single-cell validation cohorts had very few malignant cells (∼400 cells), suggesting that larger, clinically integrated single-cell EAC cohorts with sufficient malignant cells are needed to further validate our cNMF results. Fourth, the relatively small size of the ST samples necessitates caution in interpreting quantitative conclusions. Additionally, we did not generate immunofluorescence data to validate the differential spatial localization of TME programs (e.g., iCAF) or malignant programs (e.g., cNMF4), necessitating further phenotypic and spatial validation. Fifth, we did not generate matched whole genome or exome sequencing data and thus cannot exclude the impact of genetic heterogeneity on cell type and malignant program abundance. Sixth, Lastly, the heterogeneous nature of the cohort, including variations in metastatic status, treatment regimen, and anatomical location, posed a challenge.

Broadly, our study underscores the clinical importance of tumor cell heterogeneity in primary and metastatic EAC, elucidating the association of distinct tumor cell states with clinical characteristics, ICI response, and potential TME interplay, marking a key step towards understanding EAC formation and progression.

## Methods

### Experimental model and patient details

The 10 patient samples (eigh tumor tissue and two non-paired normal adjacent tissue) were collected with written informed consent and ethics approval by the Dana-Farber Cancer Institute Institutional Review Board under protocol numbers 14-408, 03-189, and 17-000. The nomenclature designates: normal adjacent tissue samples as P1 and P2; primary tissue samples as P3, P4, and P5; and metastatic samples as P6, P7, P8, P9, and P10.

### Patient tissue sample collection and dissociation for multiome snRNA-seq/ATAC-seq

Nuclei isolation was performed on frozen biopsy specimens as previously described^94^. Low-retention microcentrifuge tubes (Fisher Scientific, Hampton, NH, USA) were used throughout the procedure to minimize nuclei loss. Briefly, patient tissue was separated from optimal cutting temperature (OCT) by removing the OCT with sharp tweezers and scalpels. Tissues were then manually dissociated into a single-nuclei suspension by chopping the tissue with fine spring scissors for 10 minutes, homogenizing in TST solution, filtering through a 30 um MACS SmartStrainer (Miltenyi Biotec, Germany), and centrifuging for ten minutes at 500g at 4C. The resulting nuclei pellet was resuspended in a lysis buffer to permeabilize the nuclei before centrifuging again for 10 minutes at 500g at 4C. The final nuclei pellet was resuspended in 100 ul of 10x Genomics Diluted Nuclei Buffer and trypan blue-stained nuclei were counted by eye using INCYTO C-Chip Neubauer Improved Disposable Hemacytometers (VWR International Ltd., Radnor, PA, USA).

Approximately 16,000-25,000 nuclei per sample were loaded per channel of the Chromium Next GEM Chip J for processing on the 10x Chromium Controller (10x Genomics, Pleasanton, CA, USA) followed by transposition or cDNA generation and library construction according to manufacturer’s instructions (Chromium Next GEM Single Cell Multime ATAC + Gene Expression User Guide, Rev F). Libraries were normalized and pooled for sequencing on two NovaSeq SP-100 flow cells (Illumina, Inc., San Diego, CA, USA).

### snRNA and snATAC multiome processing

#### snRNA-seq and snATAC paired data preprocessing

The paired snRNA-seq and snATAC-seq samples were sequenced using Illumina HiSeq X. Subsequently, the raw bcl files were aligned to the human reference genome GRCh38 for each sample via Cell Ranger Arc 2.0.

#### snRNA-seq specific processing and cell type annotation

To mitigate potential ambient RNA contamination within the RNA assay of the multiome data, we used Cellbender^95^ to computationally remove ambient RNA counts from each count matrix.

After, Scrublet^96^ was employed to identify cell barcodes that may be potential doublets from the ambient RNA-adjusted RNA count matrices, and these barcodes were subsequently removed. The resulting doublet-free ambient RNA-adjusted count matrices were then employed for further downstream analyses.

RNA assay quality control procedures were conducted for each individual patient sample using Scanpy^97^. Cell barcodes with fewer than 200 unique genes expressed, genes expressed in fewer than three cells, and cell barcodes exhibiting greater than 20% of all RNA expression counts mapped to mitochondrial genes (pctMT) were filtered out. RNA expression per cell was normalized via counts per 10k (CP10k), *i.e.*, dividing the counts by the library size of the cell and normalizing to 10,000 total counts per cell, followed by log(x+1) transformation. After performing Leiden clustering (resolution = 0.7) on the 15-nearest neighbor graph of the RNA assay per individual patient sample, component cell types were manually annotated by evaluating canonical marker gene expression per cluster identified through differential expression (DE) utilizing the overestimated variance t-test.

The copy number variation (CNV) profile of each cell per individual patient sample was computed utilizing a Python implementation of InferCNV (https://github.com/icbi-lab/infercnvpy), employing a mixture of non-malignant cells as a reference (annotated fibroblasts, endothelial cells, and immune cells) based on their presence in the sample. Cells were clustered according to their CNV profile using Leiden clustering, with clusters labeled as malignant or non-malignant depending on their average CNV score. Subsequently, cells were assigned a malignant or non-malignant status based on their cluster membership per individual patient sample.

Refinement of cell type annotation was performed by analyzing cells from all patients of a single type after integration. For each major TME cell type (T/NK, myeloid, endothelial, fibroblast, muscle), cells having a relatively lower pctMT (<15%) were further analyzed downstream. We strengthened the pctMT threshold only in the TME compartment, as malignant and epithelial cells can display higher basal levels of mitochondrial counts^98^. Cells were subsetted per cell type and all cells of the same type were integrated using Harmony^99^, followed by Leiden clustering to obtain subclusters. The integration was performed on a cell-type level rather than on the full set of cells to obtain more fine-grained integration. Manual annotation of subclusters was carried out using marker genes identified through differential gene expression with an overestimated variance t-test as before.

Annotations of myeloid cell populations were cross-referenced with pan-cancer myeloid annotations from Cheng et al.^24^, while cancer-associated fibroblasts (CAF) cells were compared to pan-cancer CAF annotations from Luo *et al*.^16^. For visualization only, we integrated the fully annotated cohort using Harmony, opting not to use the cell-type-specific Harmony integration.

#### snATAC-seq specific preprocessing

The processed snATAC-seq data was acquired utilizing CellRanger Arc 2.0 (snapshot 28). Subsequently, the Signac package was employed for comprehensive processing of the ATAC data^100^ (https://stuartlab.org/signac/). Adhering to the guidelines outlined in the 10X multiome Signac vignette, the filtered counts and ATAC fragments obtained from CellRanger Arc 2.0 were utilized to re-call peaks using MACS2^101^ (https://pypi.org/project/MACS2/). Additionally, peaks located in non-standard chromosomes and genomic blacklisted regions were excluded. The consolidated peaks from all samples underwent further filtration, removing those with a width below 20 bp or exceeding 10,000 bp.

Cell type annotations were directly transferred from the snRNA annotations, as the RNA and ATAC measurements were paired. Cells excluded during standard quality control in the RNA measurements but not in the ATAC measurements were annotated as NA. Subsequently, a comprehensive quality control assessment was conducted on the entire set of cells across all samples. Cells with ATAC counts falling below 1000 or exceeding 100,000, a nucleosome signal surpassing 2, a TSS enrichment below 3, or a fraction read in peaks below 0.15 were filtered out.

Normalization of the ATAC count matrix was executed utilizing the term-frequency inverse-document-frequency (TF-IDF) transformation, following default parameters in Signac. Dimensionality reduction was carried out using Latent Semantic Indexing (LSI) with 40 components on the TF-IDF normalized matrix, with UMAP computed on the harmony-corrected LSI components.

### snRNA-seq analysis

#### Differential abundance testing

Differential abundance testing for the myeloid, CAF, and lymphoid compartments was conducted employing the milopy package ^18^ (https://github.com/emdann/milopy). Of note, the sampling bias, *i.e.*, the fact the resection from the tumor tissue and adjacent normal tissue may vary in tissue size and baseline abundance and types of cells across the tissue, as well as the processing bias, *i.e.*, the fact cells differentially suffer from dissociation and processing, might bias differential abundance testing and limit the interpretability of differential abundance results. For the myeloid compartment, differential abundance testing compared normal adjacent tissue with tumor tissue. For the CAF compartment, differential abundance testing compared primary with metastatic tissue. The Milo method was executed on the cell-type specific Harmony-corrected principal components (PC), utilizing a 20-nearest neighbors graph. Neighborhoods were assigned labels through majority voting: if over 60% of cells within a neighborhood belonged to an individual cell type, the neighborhood was labeled accordingly. Otherwise, the label “mixed” was assigned.

#### Malignant cell program discovery through consensus Negative Matrix Factorization (cNMF) and characterization

To dissect the malignant cell compartment, we employed consensus non-negative matrix factorization (cNMF)^102^ (https://github.com/dylkot/cNMF) per individual patient sample and then aggregated the results as described below. cNMF was performed on a sample-level rather than on the full cohort to avoid detecting patient-specific programs primarily driven by technical factors such as batch effects or copy-number variation (CNV) profiles. Cells annotated as putatively malignant based on canonical marker gene expression but not from clustering on inferCNVpy copy number score were filtered. For each sample, cNMF was performed on the RNA counts matrix of the 2,000 most highly variable genes, selecting the number of components (k) based on recommended criteria (i.e., inspecting the error and stability plot and picking the smallest k that minimized error while maximizing stability). Density threshold was set to 0.1 for each sample. cNMF programs expressed in too few cells or showing expression of TME-related genes, potentially indicating contamination, were manually removed.

The cNMF gene expression programs generated per individual patient sample were characterized by a vector of weights per gene representing its contribution to the program. These programs were combined across all samples by calculating their pairwise cosine similarity after removing small (high score in <10 of cells) or contaminated programs. Hierarchical clustering with an average linkage method was then applied to group similar programs into five clusters. A cNMF program was defined by the median weight of clustered gene expression programs, with the top 100 contributing genes used as a gene signature for the cNMF program. Cells from all patients were scored for the resulting cNMF gene signatures using the scanpy scoring method, i.e., the average gene expression of signature genes subtracted with the average gene expression of control genes.

The programs were compared to pan-cancer programs described in Gavish et al.^33^. For each combination of program uncovered in our dataset and program uncovered in the Gavish et al. publication, we computed the fraction of genes that were found in both programs on the number of genes from the Gavish et al. programs captured in our dataset. We also compared the programs to the Barrett’s esophagus programs described by Nowicki-Osuch et al.^1^ using Gene Set Enrichment Analysis (GSEA)^103^. Finally, GSEA^103^ was run using the prerank function on the ranked list of genes associated with each program, using the hallmarks of cancer as a search database ^104^.

#### Validation of malignant cell programs in external datasets

In order to assess the reproducibility of the malignant cell cNMF programs identified within our cohort, we conducted a similar analysis on the malignant cell compartment of two external single-cell RNA sequencing studies focusing on esophageal adenocarcinoma: the datasets from Carroll et al.^9^ and Croft et al.^10^. Following the methodology outlined in the previous section, we applied cNMF to derive programs for each sample in these external datasets. Subsequently, we computed the cosine similarity between each of these programs and the cNMF programs previously identified in our own dataset. This comparative analysis allowed us to determine the degree of recurrence and consistency of the identified programs across multiple independent datasets.

### snATAC analysis and link with snRNA

#### Link between snRNA and snATAC

To establish a connection between the programs identified in the malignant cell compartment and ATAC peaks, we calculated the Pearson correlation in malignant cells between the TF-IDF-normalized peak accessibility and program score transferred from the snRNA-seq. We then filtered out peaks in the 25% least expressed category in malignant data before performing the correlation computation. Subsequently, we determined the false-discovery rate (FDR) corrected *q*-value associated with correlation for each peak. Peaks with an FDR *q*-value below 0.05 and a Pearson correlation coefficient exceeding 0.1 were considered significantly correlated with a specific program.

#### Representative cells and link between RNA and ATAC identity

To establish the connection between the transcriptomic and epigenetic characteristics of cells, we identified representative cells for each cNMF program. Specifically, we selected cells within the top 5% highest cNMF score for each program, ensuring exclusivity by removing cells that ranked in the top 5% for two or more programs. These cells were designated as cNMF representative cells and were utilized to depict genome tracks surrounding genes of interest. Notably, due to differential recovery rates of ATAC and RNA, the proportion of representative cells with paired ATAC measurements varied.

To characterize the ATAC identity of cells, we identified the top 200 most significantly correlated regions with each cNMF program as cNMF-associated regions. Subsequently, we computed the Z-score for each region, estimating the mean and standard deviation across the population of cNMF representative cells. The ATAC data were then scored for each program using the mean Z-score of cNMF-associated regions, and each cell was assigned an ATAC identity based on the maximum score. A comparison between RNA and ATAC identities was performed using a confusion matrix.

Drawing inspiration from previous work^42^, we assigned a plasticity score to each cell using Shannon’s entropy as a measure of plasticity. We assigned a probability of belonging to a program using a softmax transformation with temperature. Let *s_j_*(*ATAC_i_*) be the ATAC score associated with cNMF_i_ in cell j; we transformed the score in probability *p_i,j_*

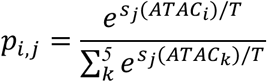

The temperature parameter T was chosen to optimize the calibration curve associated with RNA and ATAC identity correspondence (Suppl. Fig. S11). The plasticity score of cell *j* was then computed as

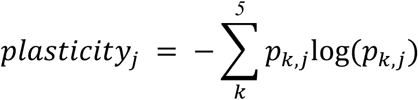

We finally computed the distribution of plasticity scores in the cNMF representative populations.

### Spatial transcriptomics (ST) analysis

#### ST data preparation and sequencing

FFPE-embedded tissue sections of 3 –10 μm thickness were sectioned then placed on a slide. H&E staining was performed by Brigham and Women’s Hospital Pathology Department core facility. When available, 2-4 FFPE scrolls of 10 – 20 μm thickness were collected in microtubes and stored at –200C. RNA quality was assessed using FFPE scrolls or from tissue sections previously placed on a slide by gently removing the FFPE section with a sterile blade and immediately transferring it to a microtube. RNA extraction was carried out using a Qiagen RNeasy® FFPE kit. RNA integrity, measured by DV200 value, was determined using the Agilent 4200 TapeStation with RNA High Sensitivity ScreenTape was used. FFPE H&E-stained slides were imaged according to the Visium CytAssist Spatial Gene Expression Imaging Guidelines Technical note. Briefly, using the Leica Aperio VERSA scanner microscope, slides were scanned at 10X magnification. Next, the hardest coverslip was removed, and the sample deparaffinized according to the 10X Genomics Visium CytAssist Spatial Tissue Preparation guide (CG000518 Rev C) and FFPE – deparaffinization and decrosslinking guide (CG000520 Rev B). Hardset coverslips were removed by immersing them in xylene for 10 minutes, twice for each slide. Then, slides were immersed in 100% ethanol for 3 minutes, 2 times, followed by immersion in 96% ethanol for 3 minutes twice and finally in 70% ethanol for 3 minutes. Slides were incubated overnight at 4^0^C before proceeding to destaining and decrosslinking according to the guidelines. Next, the slide was placed in the Visium CytAssist Tissue Slide Cassette and destained by incubating on a low profile thermocycler adapter in a thermal cycler (BioRad C1000 Touch) at 420C in 0.1 N HCL. Subsequently, decrosslinking with 10X buffers was performed at 950C for one hour. All 5 downstream steps were followed according to the Visium CytAssist Spatial Gene Expression User Guide (CB000495, Rev E) including using 6.5 mm x 6.5 mm Visium capture area slides; (1) Probe hybridization; (2) Probe ligation, (3) Probe Release & Extension; (4) Pre-amplification and SPRIselect cleanup; (5) Visium CytAssist Spatial Gene Expression – Probe Based library construction. Visium Human Transcriptome probe set v2.0 used, which contains 18,536 genes targeted by 54,5018 probes. 2.4% (451) of these genes are excluded by default due to predicted off-target activity to a different gene. All cleanup methods were performed using SPRIselect beads (Beckman Coulter), Qiagen EB buffer, and 10X Magnetic separator. Cycle number determination for GEX sample index PCR was performed using Kapa SYBR Fast qPCR Master Mix and qPCR amplification plots were visualized on the 7900HT Real-Time PCR system. Dual Index TS Set A, contains a mix of one unique i7 and one unique i5 sample index was used for sample index PCR. GEX Post-Library Construction QC was performed on Agilent TapeStation DNA High-Sensitivity ScreenTape. Libraries were normalized and pooled for sequencing on NextSeq 150 flow cells (Illumina, Inc., San Diego, CA, USA).

#### ST data preprocessing and cell type annotation

Following the spatial transcriptomics sequencing, the raw bcl files were demultiplexed using bcl2fastq and aligned to the human reference genome GRCh38 for each sample via SpaceRanger (v2.1.1). Quality control procedures were conducted individually for each patient using Squidpy^105^. Spots with fewer than 5,000 counts, genes expressed in fewer than 10 spots, and spots exhibiting over 30% reads mapped to mitochondrial DNA (pctMT) were filtered out.

The copy number variation (CNV) profile of each cell was computed utilizing a Python implementation of InferCNV (https://github.com/icbi-lab/infercnvpy). To get an initial estimate of malignant versus normal ST spots, used as input to inferCNV, we employed a method inspired from the STARCH method initialization^106^. Briefly, we ran PCA on the log(1+CP10K) normalized ST data and clustered the data using K-means (*k=2*). We assigned the cluster with the highest average expression to the tumor cluster and the remaining cluster to normal. Normal spots are used as reference for the inferCNV algorithm. We then clustered spots according to their CNV profile using Leiden clustering and assigned clusters with a strong CNV profile to tumor spots. Clusters with a similar CNV profile to the tumor spots but with a weaker overall signal were assigned to mixed spots. Finally, spots with no CNV profile or with a CNV profile opposite to the tumor profile were labeled as normal spots. Hence, this procedure yields a refined assignment to spots to mostly tumor, mixed tumor and TME, and mostly TME regions. We further refined the annotations by spatially smoothing annotations: if a tumor or normal spot contained one or zero spots in the 6-nearest neighbors of the same category, the label was reassigned to the majority label of the neighborhood (tumor, mixed, or normal).

We then computed the distance of each tumor spot to the periphery of the tumor using the shortest path to the nearest normal or mixed spot. Spots with a small assigned distance were hence located at the tumor periphery, while spots with a large assigned distance were located at the tumor core.

#### Deconvolution of ST data

To estimate the proportion of specific cell types within each spot as well as to obtain cell-type specific gene expression, we ran Cell2Location^71^ on each sample, using the full annotated snRNA-seq discovery cohort as reference. We trained the negative binomial model on the discovery cohort using default parameters to obtain estimated cell-type specific average gene profiles. We then ran the Cell2Location model, using as prior N=5 average cells per spot and alpha=20 (relaxed regularization). This yielded an estimated number of cells from a specific cell type per spot. We then sampled from the posterior distribution of the trained model to obtain cell-type specific gene expression per spot.

#### Scoring the cNMF programs and TME subtypes

We used the carcinoma-specific gene expression matrix generated by Cell2Location to score the cNMF programs, using the top 100 cNMF contributing genes as a signature, similarly as for the snRNA-seq data. The matrix was first normalized using the log(1+CP10K) transformation. The cNMF score was computed as the average expression of the Z-score of signature genes, where the Z-score of a gene is computed as 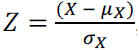, where X is the original gene expression, μ*_x_* (resp.σ*_x_*) is the gene average (resp. standard deviation) over all spots. We used a similar procedure to score the TME subtypes, using the corresponding deconvolved layer for scoring, i.e., myeloid-specific gene expression matrix for myeloid subtypes and fibroblast-specific for fibroblast subtypes.

#### Spatially constrained malignant and TME interaction with NCEM

To obtain estimates of interaction between cell types present in our data, we used node-centric expression models (NCEM)^83^ on the Cell2Location deconvolved data, using the following tutorial to prepare the data (https://github.com/theislab/ncem_benchmarks/blob/main/notebooks/data_preparation/deconvolu tion/cell2location_human_lymphnode.ipynb) and the following tutorial to process the data (https://github.com/theislab/ncem_tutorials/blob/main/tutorials/type_coupling_visium.ipynb). In brief, the intensity of the interaction is estimated as the L1 norm of the significant coefficients of the model predicting the gene expression from the cell type and niche. In the visual representation, the strength of interactions is proportional to the width of the line linking two cell types; only cell types that have more than 25 significant coefficients (FDR p<0.05) are linked in the graph.

#### Ligand-receptor interactions using CellPhoneDB

To compute the significant interactions at a local scale, we used LIGREC, a variation of CellPhoneDB implemented within Squidpy^105^, with default parameters. Given this method does not include spatial information to inform possible interactions, we constrained our analysis to spots located near the tumor periphery. We labeled tumor spots with a distance of 2 or less to the nearest tumor or mixed spot as tumor periphery, and those with a distance of 3 or more as tumor core. Normal spots with a distance of 2 or less to the nearest tumor or mixed spot were labeled as normal periphery, and those with a distance of 3 or more as normal healthy. We then ran LIGREC using these labels and restricted our analysis to significant interactions between tumor periphery and normal periphery spots. Significant interactions were hence computed only using cells located near the periphery, which does not however ensure that each spot expressing a specific ligand was in the direct periphery of the spot expressing the receptor. Although we found numerous interactions, we visually represented only significant interactions (FDR p<0.1) where the ligand or receptor belonged to the signature genes of the cNMF programs or the myeloid or fibroblast subtypes, specifically the top 100 contributing or most differentially expressed genes.

#### Enhancer-driven gene regulatory network inference

To construct an enhancer-driven gene regulatory network (GRN), we utilized the SCENIC+ software^27^. The SCENIC+ analysis was conducted at a sample level, with subsequent aggregation of the sample results. Samples with adequate ATAC recovery were included, excluding two normal adjacent samples and one primary sample (P1, P2 and P3).

#### Enhancer-driven GRN inference with SCENIC+

For each sample, we first created a pycisTopic object by integrating the filtered gene expression and cell type annotations, preprocessed according to the pipeline outlined in “snRNA-seq data preprocessing and cell type annotation,” along with the ATAC fragments obtained through the Cellranger ARC pipeline and the MACS2-called peaks. The analysis encompassed all cells that passed the Cellranger ARC filtering. Subsequently, we employed the serial Latent Dirichlet Allocation (LDA) implementation in pycisTopic, running models with 2, 4, 10, and 16 topics. The selection of the optimal model was based on a combination of metrics as recommended in the pycisTopic tutorial (https://pycistopic.readthedocs.io/en/latest/Single_sample_workflow-RTD.html).

The topic-region distributions were binarized using both the Otsu method and the top 3,000 regions per topic, as advised in the tutorial. Additionally, we computed the differentially accessible regions per cell type, utilizing the cell types annotated in the snRNA data. To identify enriched motifs in candidate enhancer regions, we executed pycisTarget with precomputed databases for motif enrichment and annotations obtained from the auxiliary data of cisTarget (https://resources.aertslab.org/cistarget/).

Subsequently, genes and regions expressed in less than 10% of the cells were filtered out, and the September 2019 Ensembl version was employed for annotation (https://sep2019.archive.ensembl.org/index.html). Finally, leveraging the paired snRNA– and snATAC-seq data and the motif enrichment matrix derived from pycisTarget, we inferred a GRN with SCENIC+ default parameters.

#### Identifying candidate master transcription factors associated with TME and malignant programs

To identify potential candidate master transcription factors (TFs) associated with cell types or programs, we analyzed the results of SCENIC+ following the tutorial (https://scenicplus.readthedocs.io/en/latest/Scenicplus_step_by_step-RTD.html). SCENIC+ provided outputs of enhancer-driven regulons (eRegulons), delineated as a transcription factor and its regulated genes and regions. The eRegulons were scored in each cell using AUCell, and the TF-eRegulon relationship was computed using pseudobulks for each cell type. High-quality eRegulons were selected, with those exhibiting a TF-eRegulon correlation below 0.2 being removed.

To identify candidate TFs associated with each major cell type in the TME, we calculated the regulon specificity score (RSS) for each eRegulon. Candidate TFs were considered associated with a TME cell type if they exhibited a significant RSS in at least two samples for this cell type. Subsequently, for each cell type, we determined the TF expression Z-score on the entire cohort, along with the associated gene-based and region-based eRegulon Z-scores. Candidate TFs were evaluated for their consistent overexpression in the cell type of interest across all three Z-scores.

To identify candidate master TFs associated with the cNMF programs, we computed the correlation for each available TF between the cNMF program scores and all three measurements of TF activity (TF gene expression, gene-based eRegulon score, and region-based eRegulon score). The TFs were ranked based on their correlation with the three measurements, and the median rank of TFs across all three measurements was computed. Only TFs with a correlation exceeding 0.1 in all three modalities were selected. The top 20 TFs with the highest correlation mean across modalities were designated as candidate TFs.

#### Validation of the identified transcription factors in external datasets

To validate the association between the inflammatory cancer-associated fibroblast (CAF) phenotype we identified and its associated transcription factors (TFs), we utilized the pan-cancer CAF atlas provided by Luo *et al*.^16^. Furthermore, to confirm the link between the revealed cNMF programs and their respective TFs, we employed two previously mentioned external single-cell validation cohorts (Croft et al.^10^ and Carroll et al.^9^).

Using the top 100 marker genes for the inflammatory CAF phenotype or the cNMF programs as signatures, we scored them using the scanpy scoring function in the external cohorts. To prevent overestimation of the correlation between the score and TF, we excluded the candidate TFs from the original signature. Due to the absence of associated scATAC-seq data, SCENIC+ could not be executed in external datasets. Therefore, we utilized the SCENIC program^107^ on the three external datasets to estimate regulon activity.

Subsequently, we computed the correlation between the inflammatory CAF score or cNMF scores and the expression of all known TFs listed by Lambert et al. ^108^ (http://humantfs.ccbr.utoronto.ca/). Additionally, we calculated the correlation between the scores and the eRegulon score as determined by SCENIC. The candidate TFs identified in our dataset through SCENIC+ were highlighted among the most highly correlated TFs, considering both their correlation with TF gene expression and eRegulon score. Of note, we only computed the correlation between the scores uncovered in Carroll *et al*. (cNMF_1_, cNMF_2_, cNMF_4_, and cNMF_5_) and Croft *et al*. (cNMF_1_, cNMF_3_, and cNMF_4_).

Furthermore, to examine whether the trajectory of candidate mTFs aligned with the hypothesized trajectories across cNMF programs, we investigated the expression of candidate mTFs linked with cNMF_4_ and cNMF_5_, which were postulated as stable opposed programs. Cells were ranked based on the difference Δ=(cNMF_4_ score – cNMF_5_ score), representing the trajectory from cNMF_5_ towards cNMF_4_. Subsequently, cells were grouped into ten equally sized bins, and the average expression level of candidate mTFs along with their associated 95% confidence interval was estimated for each bin.

#### Link between malignant programs and clinical characteristics in external bulk datasets

To ascertain whether the identified cNMF programs correlate with clinical characteristics, we assessed the scores of these programs in external bulk datasets and examined their association with various clinical parameters. To minimize the inclusion of TME components in our signature, we curated a cancer-specific signature. We selected the top 200 genes with the highest weight and subsequently filtered out genes expressed in at least 10% of any major TME cell type (endothelial, fibroblast, muscle, myeloid, lymphoid), using the remaining genes as the signature for each program.

We retrieved data from patients with esophageal adenocarcinoma from the TCGA ESCA project^7^. RNA-seq Fragments Per Kilobase of transcript per Million mapped reads (FPKM) data, non-silent mutation calls, and survival information from Liu et al. ^109^ were obtained from the UCSC Xena browser (https://xenabrowser.net/datapages/). Additionally, clinical details were directly obtained from the TCGA Network study^7^. RNA-seq expression data and clinical characteristics from the study by Hoefnagel et al.^17^ were also downloaded, with the RNA-seq raw counts transformed into transcript per million (TPM). Bulk RNA-seq expression data and associated clinical characteristics from the study by Carroll et al.^9^ were obtained and similarly transformed into TPM.

The cNMF programs were scored using single-sample Gene Set Enrichment Analysis (ssGSEA)^110^, with input data being FPKM for TCGA or TPM for Hoefnagel et al. or Carroll et al. The resulting scores were then correlated with TNM staging in the TCGA cohort. Survival analysis was conducted in the three cohorts using a univariate Cox Proportional Hazards model on standardized scores, employing default parameters from the lifelines package (https://lifelines.readthedocs.io/en/latest/fitters/regression/CoxPHFitter.html). For the Carroll et al. dataset, we used the subset of bulk expression obtained before the treatment (PreTx) to avoid introducing high correlation between patients.

#### Link between TME and malignant programs and immune checkpoint inhibitor therapy clinical benefit

To assess whether presence of specific programs, notably the inflammatory CAF program and the malignant cNMF programs, was linked to response to immune checkpoint inhibitor (ICI) therapy, we compared the distribution of score programs across patients of the Carroll et al. ^9^with or without clinical benefit. Patients were categorized into two groups based on their response to ICI therapy: those experiencing clinical benefit (CB) and those with no clinical benefit (NCB), as annotated in the Carroll et al. cohort. Notably, some patients lacked CB annotations, and patients from the operable cohort in the single-cell atlas were excluded from the CB vs. NCB comparison, participating only in patient-level comparisons. Paired measurements were obtained for patients before treatment (PreTx), after a 4-week ICI treatment window (ICI-4W), and following combined chemotherapy and ICI treatment (PostTx) when available. The distribution of inflammatory CAF and cNMF scores was compared between CB and NCB groups across PreTx and ICI-4W timepoints. Patient-level comparisons, including PostTx when applicable, were also conducted. Significance testing was performed using the Mann-Whitney U test to determine differences between scores at PreTx, ICI-4W, and PostTx measurements.

#### Ecotype analysis using BayesPrism

To explore the potential association between the identified malignant programs and the TME composition, we conducted a deconvolution analysis on the three previously described independent cohorts (TCGA^7^, Hoefnagel et al.^17^, and Carroll et al.^9^). We used the BayesPrism algorithm^111^ with default parameters, with the single-cell data from the Carroll et al. study^9^ as a reference cohort. BayesPrism provided estimates of the proportions of various cell types present in the datasets.

Inspired by the methodology outlined by Wang et al.^79^, we identified ecotypes within the datasets. Each sample was characterized based on its estimated proportion of cell types derived from the Carroll et al. dataset. Subsequently, we computed the Z-score across all samples from both cohorts regarding the proportions of cell types. Euclidean pairwise distances were then calculated between all samples, followed by hierarchical clustering with Ward linkage to group samples with similar cell type proportions.

Using the methodology described in the preceding paragraph, we scored the identified programs and assessed the enrichment of scores in the uncovered ecotypes. This analysis aimed to elucidate any potential relationships between the malignant programs and the composition of the TME across the studied cohorts.

### Prediction of drug sensitivity using scTherapy

To predict drug sensitivity for each malignant cNMF program, we employed scTherapy^73^ following its tutorial for subclone-specific drug response prediction, adapting the approach to use cNMF representative cells instead of subclones. We first identified differentially expressed genes specific to each program using these representative cells and applied the filtering criteria outlined in the tutorial to predict monotherapy sensitivity. For each program, we selected drugs predicted to have high or high-to-moderate efficacy. To identify potential combination therapies, we selected drug pairs that, together, were predicted to target all five cNMF programs.

## Supporting information

All supplementary materials

## Acknowledgements

The authors would like to thank the Single Cell Core at Harvard Medical School, Boston, MA for performing the multiome snRNA-seq/ATAC-seq sample preparation. They would also like to thank the patients and their families, as well as hospital personnel. Funding sources that supported this project include the Ambrose Monell Foundation (E.M.V.), NIH R50CA265182 (J.P.), U2CCA233195 (E.M.V.), R01CA227388 (E.M.V.), R01CA279221 (E.M.V.). T32CA092203 (C.M.A.), and Swiss National Science Foundation grant number 205321_207931 (J.Y.).

## Data availability

Raw sequencing data generated in this study are deposited in the database of Genotypes and Phenotypes (dbGaP) (https://www.ncbi.nlm.nih.gov/gap/) with accession number phs003438.v1. Existing single-cell and bulk sequencing data can be downloaded from EGA (EGAS00001006468, EGAS00001006469) for the Carroll et al. paper. Remaining existing single-cell data can be downloaded from the Gene Expression Omnibus (GEO) website, accession numbers GSE222078 (Croft *et al.*) and GSE210347 (Luo *et al.*). Remaining existing bulk data can be downloaded from the GEO website, accession number GSE207527, for the Hoefnagel *et al.* data, and from the UCSC Xena browser for the Cancer Genome Atlas ESCA cohort (https://xenabrowser.net/datapages/?cohort=TCGA%20Esophageal%20Cancer%20(ESCA)&removeHub=https%3A%2F%2Fxena.treehouse.gi.ucsc.edu%3A443). Details on how to download the data used for the analysis is detailed on Github: https://github.com/vanallenlab/EAC-multiome. Source data are provided with this paper.

## Code availability

All the code needed to reproduce this analysis is available on Github at the following address: https://github.com/vanallenlab/EAC-multiome.

During the preparation of this work the author(s) used ChatGPT to correct language and reformulate. After using this tool/service, the author(s) reviewed and edited the content as needed and take(s) full responsibility for the content of the publication.

## Ethics declaration

Declarations of interest

E.M.V.:

Advisory/Consulting: Enara Bio, Manifold Bio, Monte Rosa, Novartis Institute for Biomedical Research, Serinus Bio, TracerDx

Research support: Novartis, BMS, Sanofi, NextPoint

Equity: Tango Therapeutics, Genome Medical, Genomic Life, Enara Bio, Manifold Bio, Microsoft, Monte Rosa, Riva Therapeutics, Serinus Bio, Syapse, TracerDx

Travel reimbursement: None

Patents: Institutional patents filed on chromatin mutations and immunotherapy response, and methods for clinical interpretation; intermittent legal consulting on patents for Foaley & Hoag Editorial Boards: *Science Advances*

A.J.A. has consulted for Anji Pharmaceuticals, Affini-T Therapeutics, Arrakis Therapeutics, AstraZeneca, Boehringer Ingelheim, Kestrel Therapeutics, Merck & Co., Inc., Mirati Therapeutics, Nimbus Therapeutics, Oncorus, Inc., Plexium, Quanta Therapeutics, Revolution Medicines, Reactive Biosciences, Riva Therapeutics, Servier Pharmaceuticals, Syros Pharmaceuticals, T-knife Therapeutics, Third Rock Ventures, and Ventus Therapeutics. A.J.A. holds equity in Riva Therapeutics and Kestrel Therapeutics. A.J.A. has research funding from Amgen, AstraZeneca, Boehringer Ingelheim, Bristol Myers Squibb, Deerfield, Inc., Eli Lilly, Mirati Therapeutics, Nimbus Therapeutics, Novartis, Novo Ventures, Revolution Medicines, and Syros Pharmaceuticals.

## FIG

