## Supplementary material for "Cell states and neighborhoods in distinct clinical stages of primary and metastatic esophageal adenocarcinoma": All supplementary materials

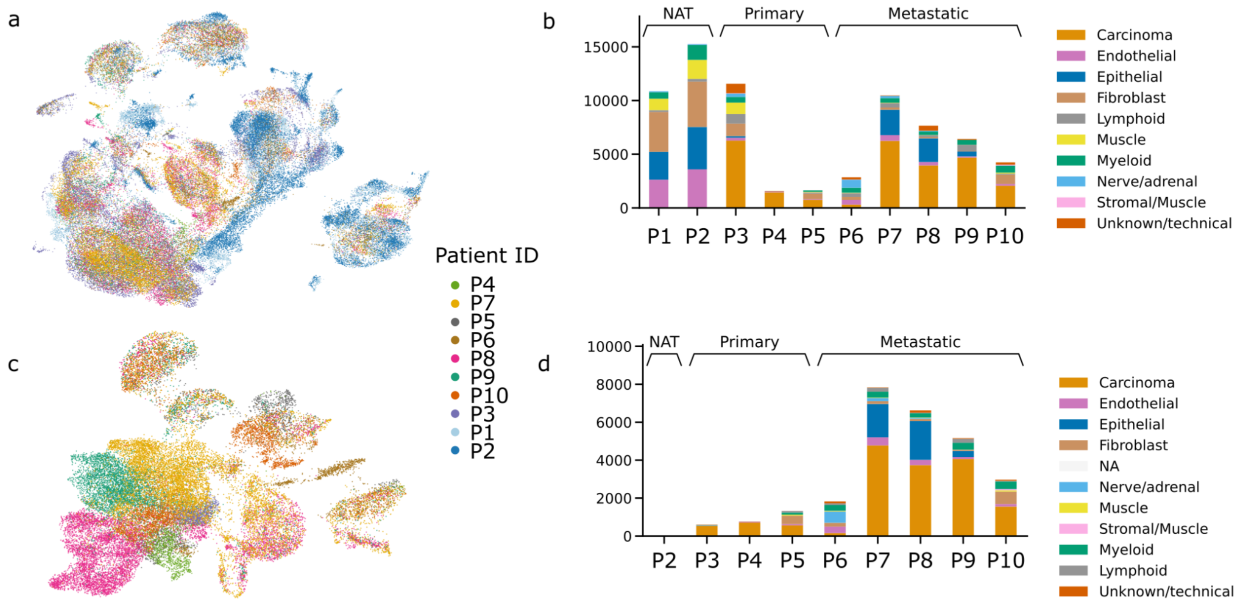


**Supplementary Figure 1: EAC primary and metastatic samples show a diverse landscape of tumor microenvironment and malignant cells in transcriptomic and epigenetic space. a,** UMAP representation of the full cohort in Harmony-corrected integrated transcriptomic data. Cells are colored according to patient ID. **b,** Counts of annotated cell types per patient for transcriptomic data. **c,** UMAP representation of the full cohort in Harmony-corrected integrated ATAC data. Cells are colored according to patient ID. **d,** Counts of annotated cell types per patient for ATAC data.

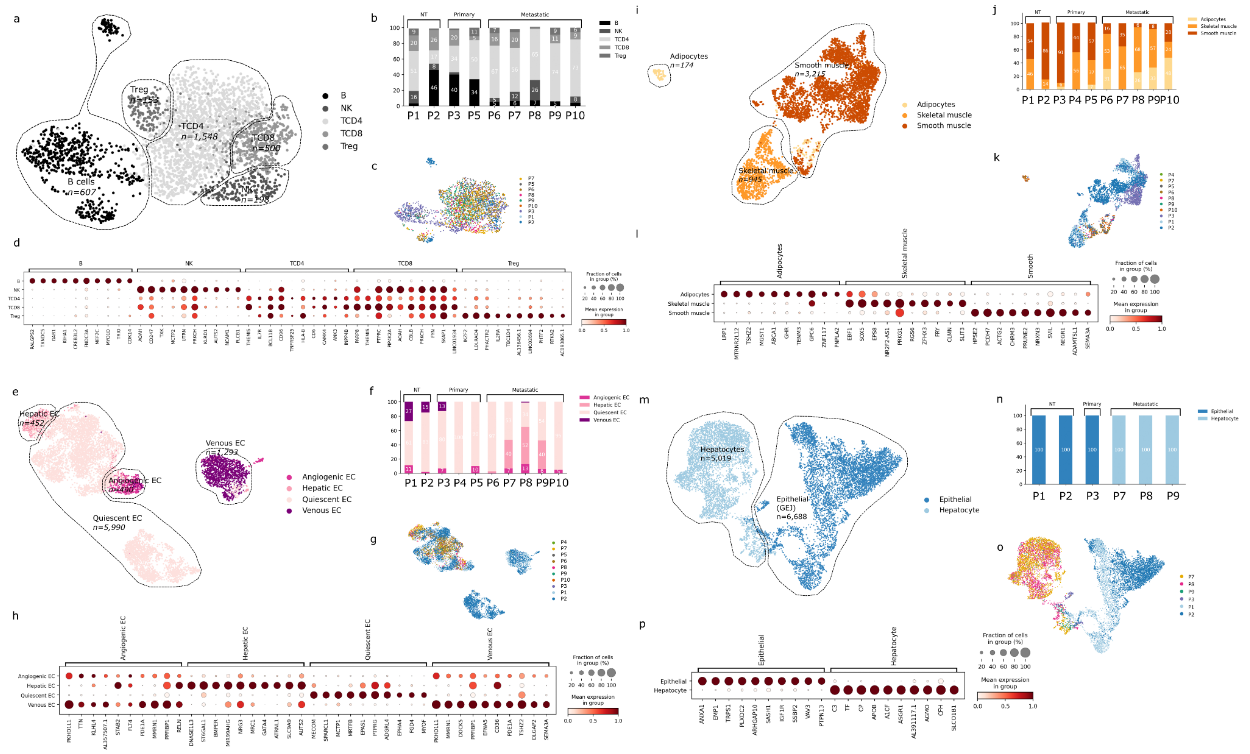

**Supplementary Figure 2: Tumor microenvironment (TME) compartments of EAC cohort. a**, UMAP representation of the lymphoid compartment in harmony-corrected integrated transcriptomic space. The annotated subtypes are indicated. **b,** Proportion of lymphoid cell types per patient. **c**, UMAP representation of the lymphoid compartment annotated according to sample of origin. **d**, Marker genes of annotated lymphoid cell types. The markers are found by running differential gene expression between cells in the compartment. Cells are grouped according to their subtype and the proportion of cells expressing the marker gene as well as the mean expression are indicated.

**e**, UMAP representation of the endothelial compartment in harmony-corrected integrated transcriptomic space. The annotated subtypes are indicated. **f**, Proportion of endothelial cell types per patient. **g**, UMAP representation of the endothelial compartment annotated according to sample of origin. **h**, Marker genes of annotated endothelial cell types. The markers are found by running differential gene expression between cells in the compartment. Cells are grouped according to their subtype and the proportion of cells expressing the marker gene as well as the mean expression are indicated. **i**, UMAP representation of the muscle compartment in harmony-corrected integrated transcriptomic space. The annotated subtypes are indicated. **j**, Proportion of muscle cell types per patient. **k**, UMAP representation of the muscle compartment annotated according to sample of origin. **l**, Marker genes of annotated muscle cell types. The markers are found by running differential gene expression between cells in the compartment. Cells are grouped according to their subtype and the proportion of cells expressing the marker gene as well as the mean expression are indicated. **m**, UMAP representation of the normal epithelial compartment in harmony-corrected integrated transcriptomic space. The annotated subtypes are indicated. **n**, Proportion of normal epithelial cell types per patient. **o**, UMAP representation of the normal epithelial compartment annotated according to sample of origin. **p**, Marker genes of annotated normal epithelial cell types. The markers are found by running differential gene expression between cells in the compartment. Cells are grouped according to their subtype and the proportion of cells expressing the marker gene as well as the mean expression are indicated.

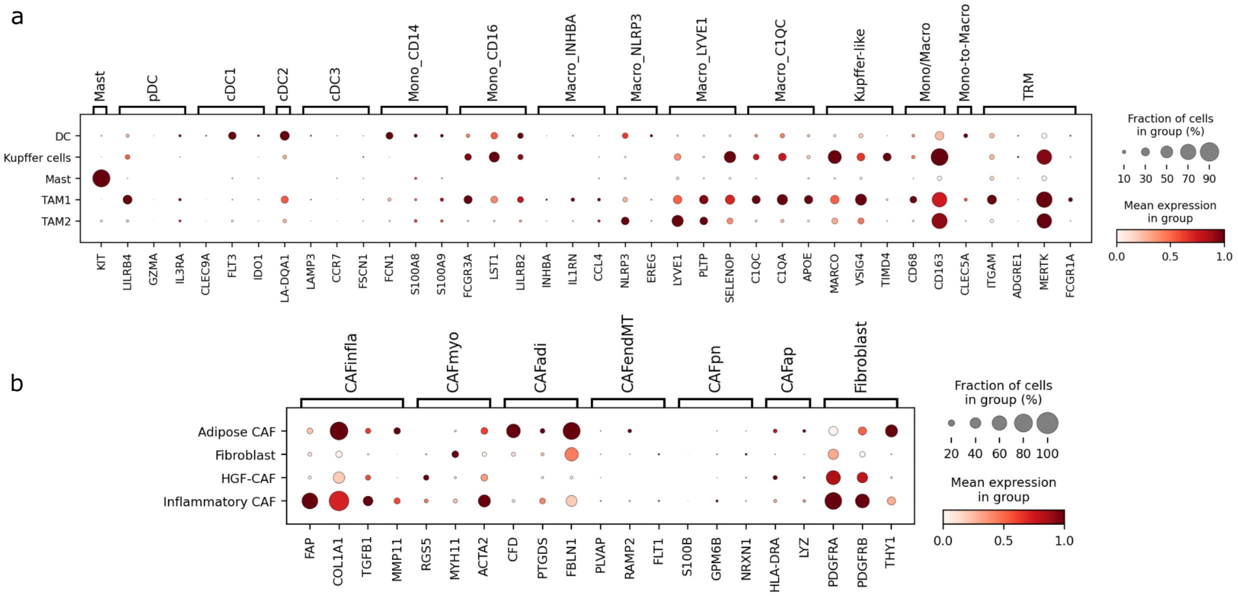

**Supplementary Figure 3: Expression of known marker genes. a**, For each annotated myeloid cell type, the fraction of cells in the group expressing marker genes from a pan-cancer myeloid study[1] and other known marker genes is represented as well as their mean expression within the group. **b**, For each annotated cancer associated fibroblast (CAF) type, the fraction of cells in the group expressing marker genes from a pan-cancer CAF study [2] and other known marker genes is represented as well as their mean expression within the group.

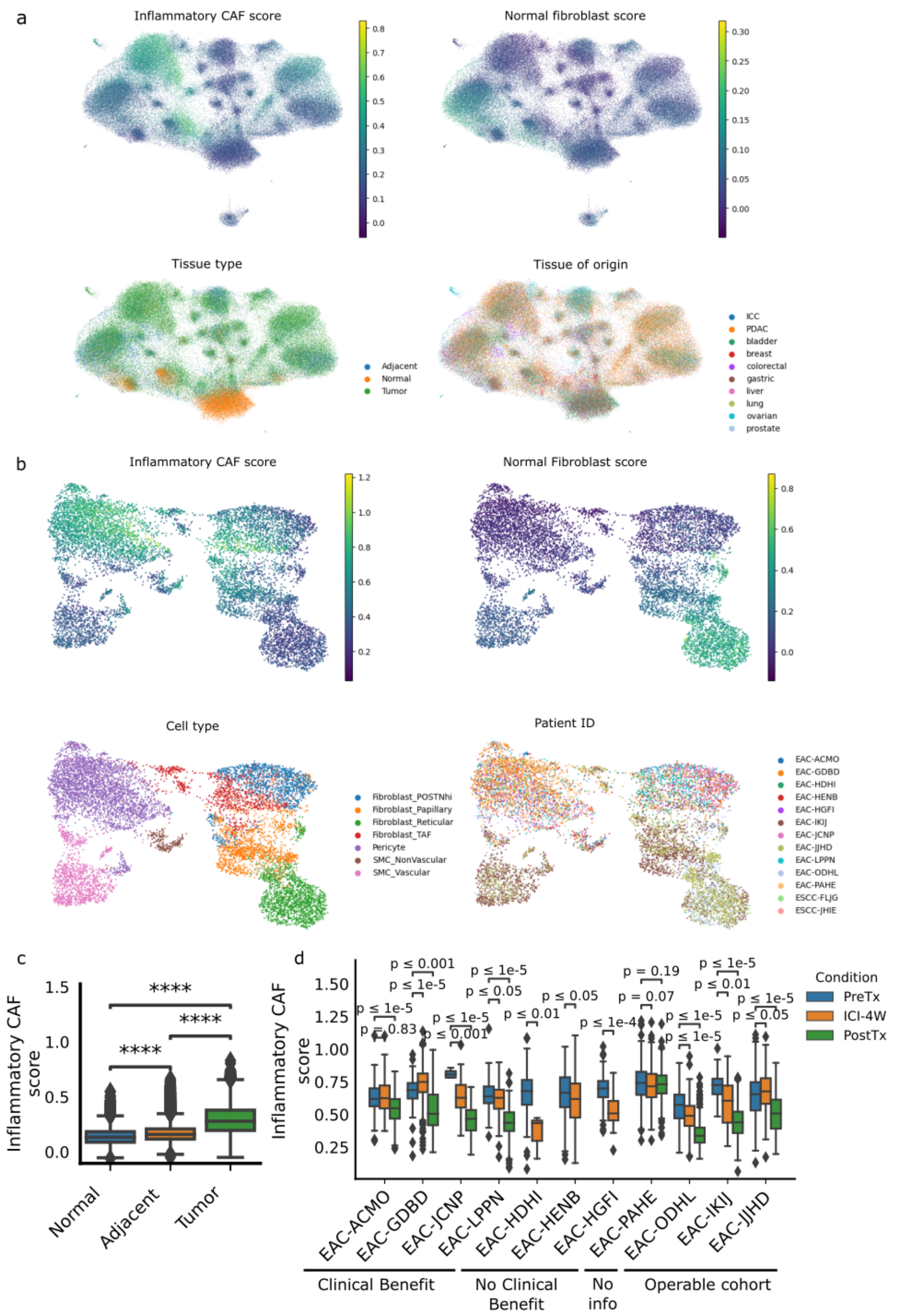


**Supplementary Figure 4: Inflammatory CAF compartment in external cohorts. a,** Harmony-corrected UMAP representation of the pan-cancer fibroblast atlas [2], highlighted according to the inflammatory CAF score and normal fibroblast score derived in our study, as well as the tissue type (normal, normal adjacent to tumor, and tumor) and the tissue of origin. **b,** Harmony-corrected UMAP representation of the Carroll et al. [3] cohort stromal compartment, highlighted according to the inflammatory CAF score and normal fibroblast score derived in our study, as well as the cell type and the patient of origin. **c,** Distribution of the inflammatory CAF score in the pan-cancer CAF atlas across tissue types. **d**, Inflammatory CAF score distribution per patient in the Carroll et al. [3] cohort. The Inflammatory CAF program is scored on the full cohort. Distribution of the score in the stromal compartment is plotted for all patients broken down according to sampling time of the biopsy (pre-treatment PreTx, after 4 weeks of ICI therapy ICI-4W, and after addition of chemotherapy PostTx). Patients are grouped according to their response to ICI therapy: clinical benefit, no clinical benefit, or no information. Distributions of scores are also indicated for patients of the operable cohort that were added to the single-cell atlas. Significance testing is performed using the Wilcoxon rank-sum test for all comparisons.

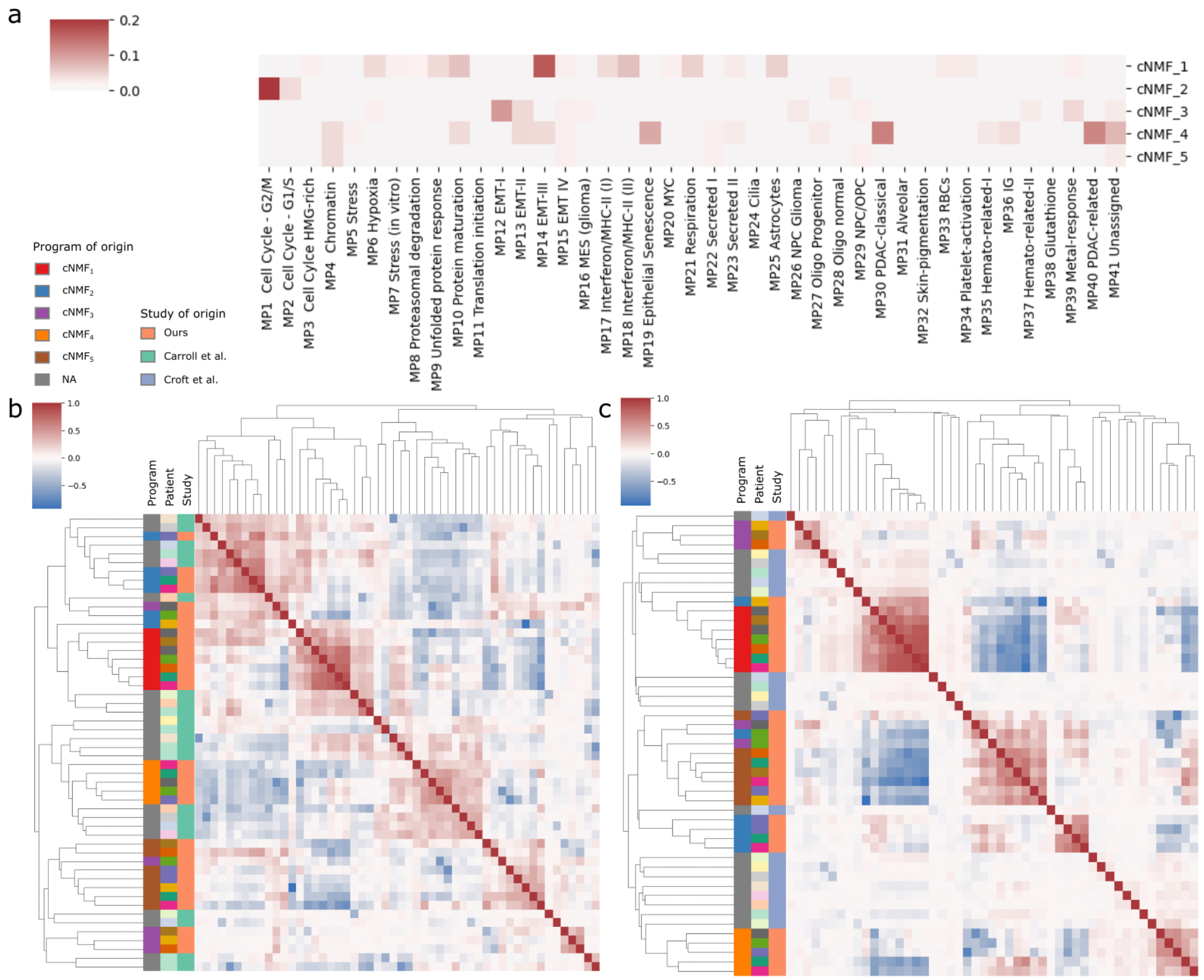


**Supplementary Figure 5: Characterization of cNMF programs linked to external studies. a,** Fraction of overlap between signature genes from cNMF programs uncovered in our study and gene programs uncovered in the Gavish et al. pan-cancer study[4]. For each cNMF program and Gavish et al. gene program, we select the top 50 genes as signature genes and then compute the fraction of shared genes between two programs as a measure of similarity. **b-c,** Clustered heatmap of the programs derived from our study and b, from the Carroll et al. study[3] and c, the Croft et al. study[5]. The pairwise cosine similarity between all programs is computed and used as a metric for hierarchical clustering with average linkage.

The first column represents the assigned program (colored for those discovered in our study or in gray in the validation cohort), the patient of origin, and the study of origin.

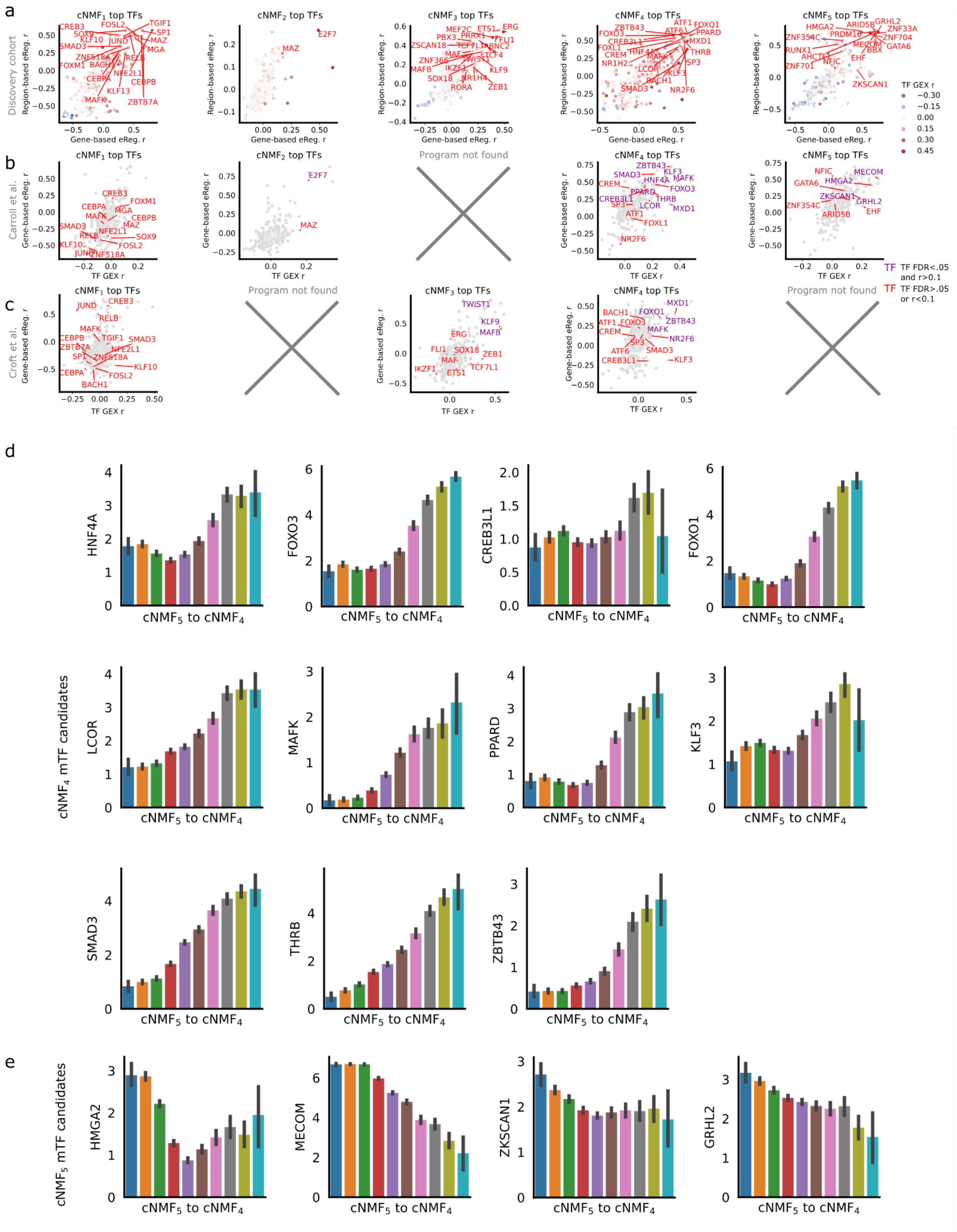

**Supplementary Figure 6: Regulatory landscape of EAC malignant programs and identification of candidate master transcription factors. a,** Candidate master transcription factors (mTFs) identified by the SCENIC+ algorithm in our cohort for all programs. The x-axis represents the correlation between the score and the gene-based eRegulon Area Under the Curve (AUC) Z-score, the y-axis represents the correlation between the score and the region-based eRegulon AUC Z-score, and the hue represents the correlation between the score and TF gene expression. Selected TFs include the top 20 TFs with the highest median rank in terms of correlation across the three measurements, correlated with an r>0.1 in each modality. **b-c,** Correlation of all available TFs' gene expression and SCENIC-estimated gene-based eRegulon score with cNMF scores in the validation cohorts of Carroll *et al.* [3] (b) and Croft *et al.* [5] (c). Candidate mTFs identified in our cohort are annotated. TFs correlated with Pearson's R>0.1 in both modalities in the external cohort are annotated in purple, while other candidate TFs are annotated in red. Programs not uncovered in the validation cohort are indicated by a cross. **d-e,** Evolution of candidate mTF expression along the $cNMF_4$ to $cNMF_5$ axis for **d,** $cNMF_4$ mTF candidates and **e,** $cNMF_5$ mTF candidates. The cells are scored using the $cNMF_4$ and $cNMF_5$ signatures. They are then broken into 10 equally sized bins according to the difference between their $cNMF_4$ and $cNMF_5$ scores, where cells with a high $cNMF_4$ and low $cNMF_5$ score are in the last bin. The expression of candidate mTFs are plotted according to the bins with standard error indicated as a bar.

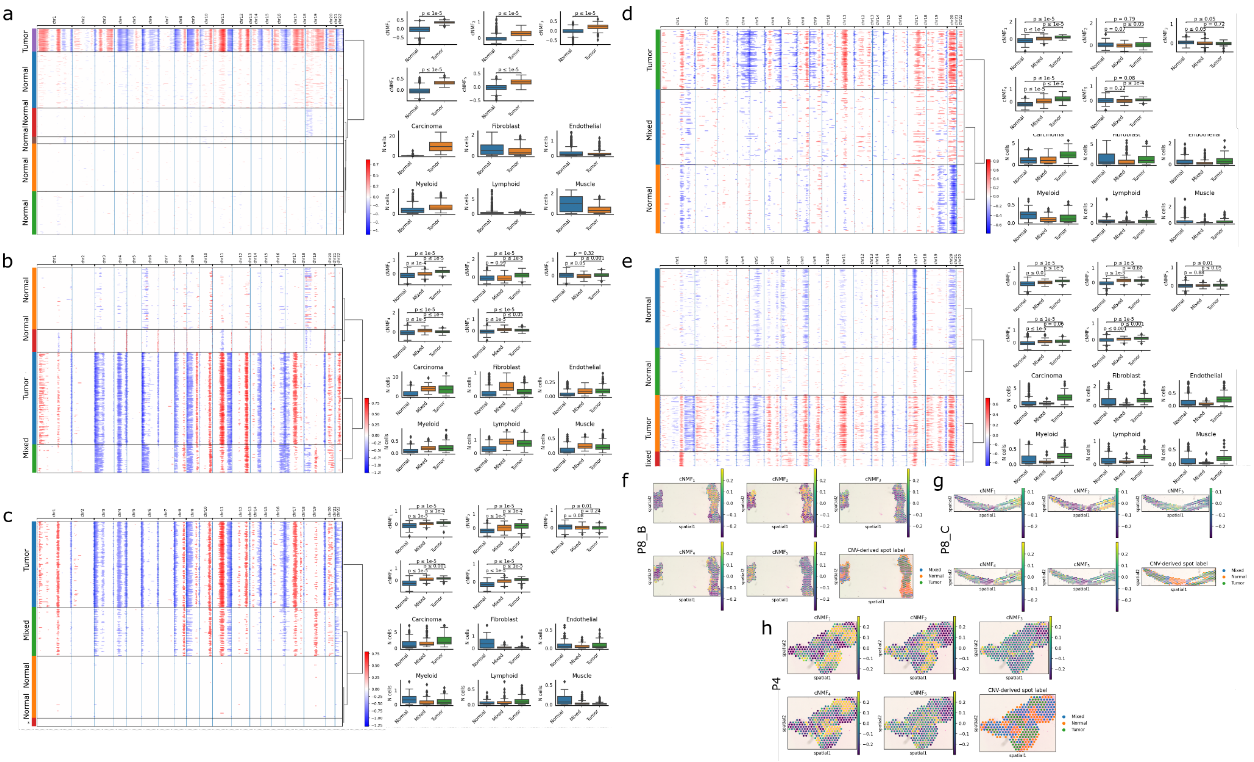

**Supplementary Figure 7: Spatial transcriptomics analysis of a subset of discovery cohort samples. a-e,** Clusters of inferCNV-derived (https://infercnvpy.readthedocs.io/en/latest/index.html) CNV profile of spots, assigned to tumor, mixed, or normal status; distribution of cNMF scores between normal, mixed, and tumor spots; and Cell2Location [6] estimated number of cells of the 6 major compartments between normal, mixed, and tumors spots, for **a,** P8 primary tumor A, **b,** P8 primary tumor B, **c,** P8 liver metastasis tumor C, **d,** P4 primary tumor, and **e,** P5 primary tumor. **f-h,** Spatial transcriptomics (ST)

slides of **f,** P8 primary tumor B, **g,** P8 metastatic tumor C, and **h,** P4 primary tumor, colored according to cNMF program score and the CNV-derived label. For each spot, we infer the CNV profile with inferCNV and assign spots to tumor, mixed, and normal status. cNMF scores are computed as the average Z-score of signature genes using the deconvolved carcinoma-specific gene expression profile of spots derived with Cell2Location[6].

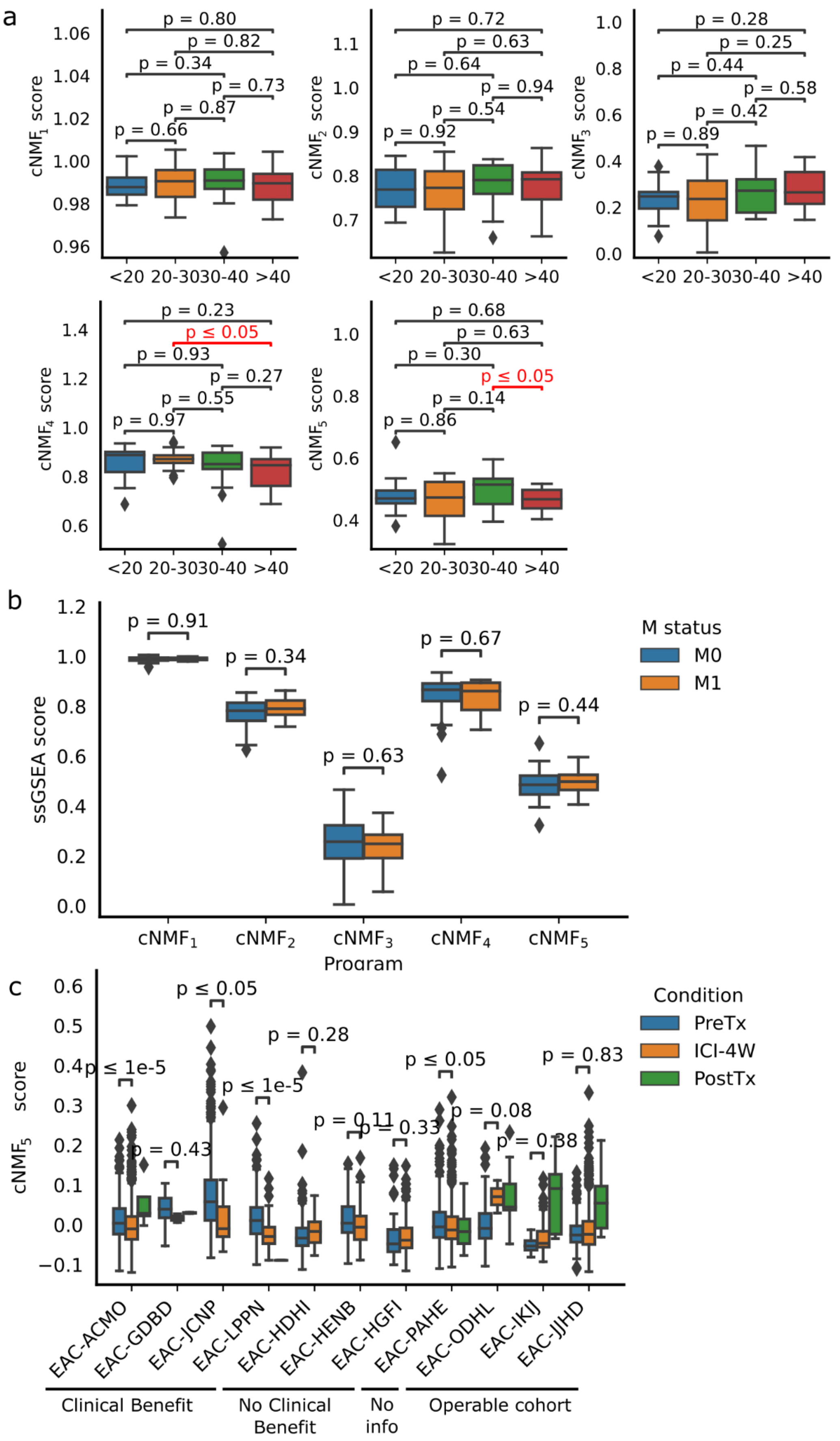
a
p = 0.80
p = 0.82
p = 0.34
p = 0.73
p = 0.87
p = 0.66
cNMF1 score
<20 20-3030-40 >40
p = 0.72
p = 0.63
p = 0.64
p = 0.94
p = 0.54
p = 0.92
cNMF2 score
p = 0.28
p = 0.25
p = 0.44
p = 0.58
p = 0.42
p = 0.89
cNMF3 score
p = 0.23
p ≤ 0.05
p = 0.93
p = 0.27
p = 0.55
p = 0.97
cNMF4 score
p = 0.68
p = 0.63
p = 0.30
p ≤ 0.05
p = 0.14
p = 0.86
cNMF5 score
b
p = 0.91
p = 0.34
p = 0.63
p = 0.67
p = 0.44
ssGSEA score
M status
M0
M1
cNMF1
cNMF2
cNMF3
cNMF4
cNMF5
Program
c
p ≤ 1e-5
p = 0.43
p ≤ 0.05
p ≤ 1e-5
p = 0.28
p = 0.11
p = 0.33
p ≤ 0.05
p = 0.08
p = 0.38
p = 0.83
cNMF5 score
Condition
PreTx
ICI-4W
PostTx
EAC-ACMO
EAC-GDBD
EAC-JCNP
EAC-LPPN
EAC-HDHI
EAC-HENB
EAC-HGFI
EAC-PAHE
EAC-ODHL
EAC-IKIJ
EAC-JJHD
Clinical Benefit
No Clinical Benefit
No info
Operable cohort

**Supplementary Figure 8: Link between cNMF scores and clinical and molecular characteristics. a,** Link between cNMF scores and homologous recombination deficiency score (HRD score) in the TCGA cohort [7]. The HRD score is downloaded from the supplementary information of the associated paper; patients are then separated into 4 bins according to their score (<20, 20-30, 30-40, >40). **b,** Link between cNMF scores and M status in the TCGA cohort [7]. **c**, $cNMF_5$ score distribution per patient in the Carroll et al. [3] cohort. The $cNMF_5$ program is scored on the full cohort. Distribution of the score in the malignant compartment is plotted for all patients broken down according to sampling time of the biopsy (pre-treatment PreTx, after 4 weeks of ICI therapy ICI-4W, and after addition of chemotherapy PostTx). Patients are grouped according to their response to ICI therapy: clinical benefit, no clinical benefit, or no information. Distributions of scores are also indicated for patients of the operable cohort that were added to the single-cell atlas. Significance testing is performed using the Wilcoxon rank-sum test for all comparisons.

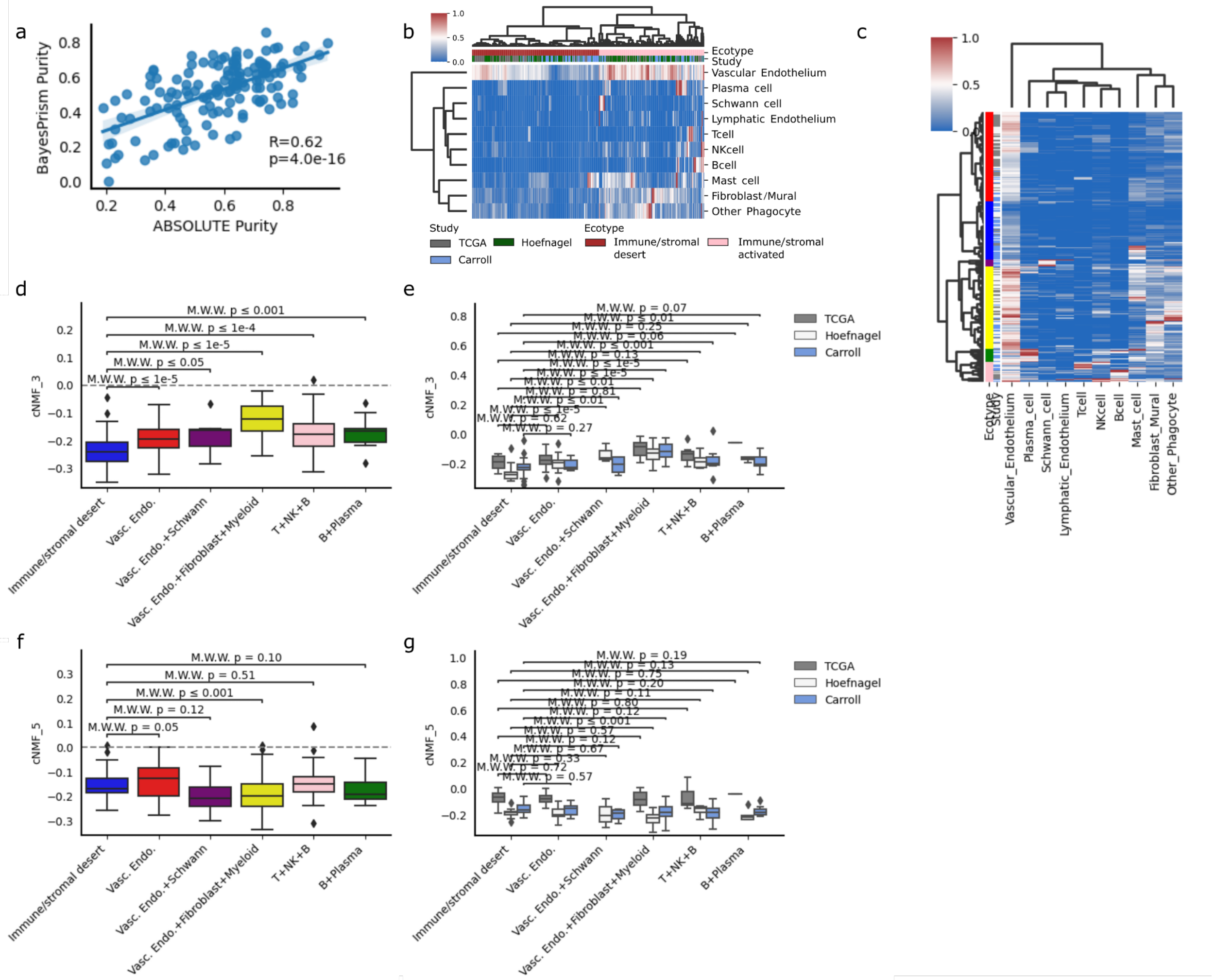


**Supplementary Figure 9: Ecotype analysis using BayesPrism and link with malignant programs. a,** Correlation between BayesPrism estimated purity and ABSOLUTE-estimated purity in the TCGA cohort. **b-c,** Ecotype analysis of the deconvolved BayesPrism data for TCGA, Hoefnagel et al., and Carroll et al.

cohorts. TME proportions are clustered using hierarchical clustering with Ward linkage. The uncovered ecotypes and study of origin are indicated, for **b,** two ecotypes and **c**, 6 ecotypes. **d-g,** Distribution of **d-e,** $cNMF_3$ scores and **f-g,** $cNMF_4$ across ecotypes. **d** and **f** represent distributions across all studies while **e** and **g** represent distributions per study. Statistical significance is computed using the Wilcoxon rank-sum test for all comparisons.

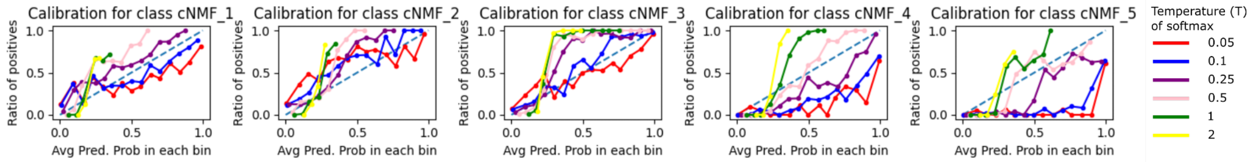


**Supplementary Figure 10: Calibration curves associated with classification of cNMF representative cells to cNMF ATAC identity for different temperature parameters for softmax transformation.** Each cNMF representative cell is assigned an ATAC cNMF score by averaging the Z-score of cNMF-associated regions. The scores are transformed into probabilities using a softmax transformation with varying temperature parameters. The calibration curve is computed by binning observations according to their predicted probability of being in the cNMF class in question. The ratio of observations truly from this class is then computed in each bin. In a perfectly calibrated model, these ratios are equal.
